# Context transcription factors establish cooperative environments and mediate enhancer communication

**DOI:** 10.1101/2023.05.05.539543

**Authors:** Judith F. Kribelbauer, Olga Pushkarev, Vincent Gardeux, Julie Russeil, Guido van Mierlo, Bart Deplancke

**Affiliations:** Laboratory of Systems Biology and Genetics, Institute of Bioengineering, School of Life Sciences, École Polytechnique Fédérale de Lausanne (EPFL), Lausanne, Switzerland; Swiss Institute of Bioinformatics, Lausanne, Switzerland

## Abstract

Many enhancers play a crucial role in regulating gene expression by assembling regulatory factor (RF) clusters, also referred to as condensates. This process is essential for facilitating enhancer communication and establishing cellular identity. However, how DNA sequence and transcription factor (TF) binding instruct the formation of such high RF environments is still poorly understood. To address this, we developed a novel approach leveraging enhancer-centric chromatin accessibility quantitative trait loci (caQTLs) to nominate RF clusters genome-wide. By analyzing TF binding signatures within the context of caQTLs, we discovered a new class of TFs that specifically contributes to establishing cooperative environments. These “context-only” TFs bind promiscuously with cell type-specific pioneers, recruit coactivators, and, like super enhancers, render downstream gene expression sensitive to condensate-disrupting molecules. We further demonstrate that joint context-only and pioneer TF binding explains enhancer compatibility and provides a mechanistic rationale for how a loose TF syntax can still confer regulatory specificity.

## Introduction

An important step in gene regulation is the recruitment of transcription factors (TFs) to enhancers. TF recruitment relies on the sequence-specific recognition of target sites by the TF’s DNA binding domain (DBD). Binding specificities for the majority of human TFs, have now been mapped^1^ and are accessible in the form of scoring matrices via several repositories^2–5^. Individual TF motif matches alone, however, are poor predictors for *in vivo* binding, with the majority of potential binding sites left unoccupied^6^. Competition with nucleosomes, sensitivity to DNA modifications^7^, recruitment of transcriptional co-activators and -repressors^8^, TF complex formation^9^, as well as interactions with RNAs^10^, all contribute to TF occupancy^11^.

To account for the influence of genomic context, deep neural network models^12,13^ have been designed to detect dependencies between motif instances, and even interactions among regulatory element (REs) dozens of kilobases apart^14^. Although such models quite accurately predict TF binding and chromatin states, even the most advanced ones do not generalize well to unseen contexts^14^, nor do they capture enhancer-based gene regulation particularly well^15,16^. Progress is hampered by the difficulty to distill human interpretable, and generalizable mechanisms from existing models. The emerging consensus is that context specificity is driven by ‘soft’ motif syntax rules^12,13^ with fuzzy molecular underpinnings. Soft syntax may thereby be a direct consequence of the quantitative nature of TF binding^17^, where occupancy and downstream regulatory output is determined by the sequence specificity and the local concentration of TFs^18–21^.

Imaging experiments have indeed shown that TFs segregate into hubs with high localized concentration within the nucleus^18,22–25^. These assemblies are referred to as condensates or regulatory factor (RF) clusters and incorporate various coactivators^26,27^, such as Bromodomain-containing proteins (BRDs). For several loci, RF clusters have been shown to form at enhancers^23–25^, in particular Super Enhancers (SEs) ^26^ that are defined by a high level of clustered regulatory activity^29,30^. However, it is still poorly understood what drives the formation of condensates at enhancers ^31^. One contributing factor is communication with the epigenomic state, e.g. the interaction of BRD4 with H3K27 acetyl lysine residues^28^. More generally, condensate formation involves weak protein-protein interactions driven by intrinsically disordered domains that are present in both cofactors and TFs^27,32,33^. Finally, TF binding to DNA itself is critical as it catalyzes condensate formation^34,35^ and directs the localization of RF clusters within cells^36^. Although there are several well-studied examples that underscore the importance of RF cluster formation and their subsequent size control for gene regulation^28,37,38^, it has proven difficult to disentangle which enhancers will give rise to RF clusters and which ones won’t. The main reason for this shortcoming is a lack of data that reliably links condensates to enhancers genome wide. Indeed, imaging approaches are limited by the number of RFs and loci that can be visualized at once and bulk epigenomic profiling cannot distinguish between signal created through strong, but independent binding of TFs or through RF cluster formation.

To overcome these limitations, we considered the biophysical properties underlying condensate formation: condensates rely on cooperativity among RFs, which leads to nonlinear behavior when RF concentration reaches a critical level^35,38,39^. The transition from an ‘independent binding’ to a high RF-concentration state occurs abruptly, meaning that slight increases (e.g., the addition of one extra molecule) can lead to large changes in subsequent RF recruitment^39^. In the context of an enhancer, such a ‘seed’ event may be mediated by the binding of one additional TF. We thus reasoned that in order to uncover enhancer-linked RF clusters, we should focus on enhancers where the addition of a single TF binding event leads to a large change in overall RF occupancy. This scenario can in fact be found at enhancer-centric chromatin accessibility quantitative trait loci (caQTLs)^40–42^ that contain a single nucleotide polymorphisms (SNPs) that creates a new TF binding site which in turn leads to a large overall change in DNA accessibility.

caQTL mapping can also aid in uncovering RE communication^42,43^, which can be interpreted as the merging of two or more compatible RF clusters – a hallmark of condensates^39^. Specifically, communication can be inferred whenever the epigenomic signatures of two or more REs covary in function of the underlying genotype. An example thereof is the *AXIN2* locus, where the creation of a single TF binding site within a TSS-proximal enhancer leads to a wide-spread, coordinated activation of multiple REs over a genomic region of >100kb^44^. For simplicity, we refer to such genetically encoded RE coordination as chromatin modules (CMs)^45^.

Here, by leveraging available data on caQTLs and CMs in lymphoblastoid cells, we uncover a new class of TFs whose dedicated function appears linked to the establishment of cooperative environments. Although these ‘context-only’ TFs are not associated with DNA accessibility directly, their binding sites nonetheless occur alongside those of cell-type specific pioneer TFs, with whom they engage in a promiscuous motif syntax. In a series of computational and experimental validation analyses, we show that the combined binding and function of both classes of TFs is consistent with the formation of RF clusters at the majority of caQTL enhancers.

## RESULTS

### Uncovering TF signatures that create caQTL environments

We used existing, fine-mapped^41^ caQTL data obtained across 100 human lymphoblastoid cell lines (LCL) measured using ATAC-seq^43^ to nominate putative RF clusters at enhancers, defined as distal or intronic regions that significantly change peak accessibility due to an underlying SNP (**Figure 1a** & **Supplemental Figure 1a,b).** We refer to these REs as ‘caQTL enhancers’. As a control, we chose enhancers that contain SNPs without significant accessibility changes, or ‘no-effect’ enhancers (**Figure 1a**). To minimize confounders, we defined peaks of equal length and matched enhancers in terms of GC content and the SNP to the peak center distance (**Figure 1b** and **Supplemental Figure 1c**). This resulted in roughly 9500 enhancers that are equally split across the two groups (i.e. ‘caQTL’ vs. ‘no-effect’ enhancers). For robustness, we created a total of fifty matched controls for caQTL enhancers (**see Methods).** Plotting the effect size of each SNP against overall enhancer accessibility in GM12878, the LCL frequently used as a reference (**Figure 1c**), we found that caQTL enhancers are more accessible than no-effect ones – a fact that arises from the count statistics used for caQTL inference (the higher the baseline coverage, the smaller the effect size required for significance) (**Figure 1c**). To account for this and other possible confounders, we assessed three separate aspects of TF function (**Figure 1d**): i) the TF’s overall association with DNA accessibility (accessibility score), which we derive using randomly sampled enhancers independent of caQTL status; ii) the TF’s ability to initiate (pioneer) an increase in accessibility at the caQTL SNP itself (initiator score); and iii) the TF’s importance in the sequence context of caQTL SNPs (context score; **Supplemental Figure 1e)**. The context score specifically interrogates whether a group of TFs that are distinct from initiators may contribute to the creation of cooperative environments. When required, we controlled for sequence composition bias and baseline accessibility (**see Methods**). We also assessed a TF’s ability to act as a repressor (inverse of the initiator score). We did not pursue it further as only a minority of TFs showed evidence for repressive mechanisms (**Supplemental Figure 1d**).

**Figure 1.**
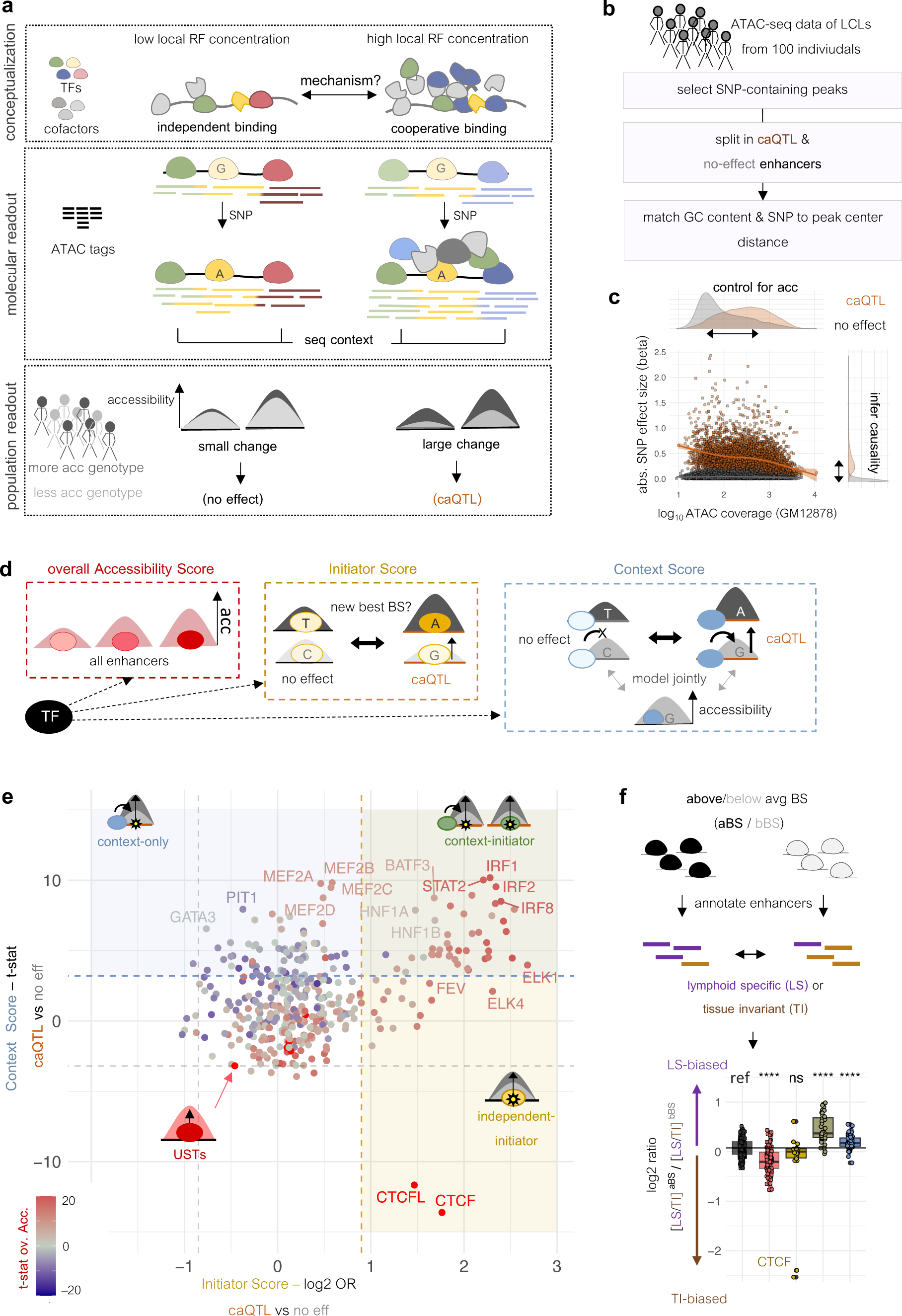
Enhancers rely on distinct classes of TFs to create caQTL environments. **a** Schematic describing TF binding under different RF factor concentrations, with low concentrations a result of independent binding and high concentrations of cooperative binding. When SNPs impact TF binding, we expect context-dependent differences in molecular readouts, highlighted both at the level of individual ATAC-seq tags (middle) and enhancer-wide accessibility changes using population genetics (bottom). **b** Process to prioritize SNPs based on whether they impact enhancer-wide accessibility in cis or not. **c** Comparison of absolute (abs.) SNP effect sizes against baseline enhancer accessibility in GM12878 (LCL reference). A negative relationship between the two measures for caQTLs results in a bias towards enhancers with higher baseline accessibility. **d** Schematic describing three ways of assessing TF function based on DNA accessibility information. First, relating binding site strength directly to accessibility using randomly sampled enhancers (left). Second, comparing caQTLs to no-effect SNPs in terms of how frequently SNPs create a new best binding site in an enhancer (middle), and third, relating binding site strength in the sequence contexts of SNPs to the SNP effect size status, while controlling for baseline enhancer accessibility (right). **e** TF context score (y-axis) versus initiator score (x-axis). Each point represents a specific TF. Accessibility scores are overlayed with a blue-red color gradient. Colored boxes indicate three of the four distinct TF classes: context-only (blue), context-initiator (green) and independent-initiator (yellow). The scatter of points in the ‘no-effect’ direction is used to define thresholds (grey and colored dotted lines). P-value cutoff for USTs is not shown but tracks with the intensity of the red color scale. **f** TF specificity towards ‘lymphoid-specific’ (LS) and ‘tissue-invariant’ (TI) elements using the DHS index. LS/TI ratios are compared across enhancers split by above or below average binding site strength (aBS,bBS) for a given TF. Y-axis shows the log2 ratio of ratios (LI/TI in aBS verus bBS).

### TFs can be stratified into four classes with varying relevance for the creation of caQTL environments

Comparing TFs based on the three scores (**Figure 1e),** we identified distinct groupings, each with different associations with caQTL-environments. Among TFs with a high initiator score are TFs identified in the original caQTL study^43^, including PU.1, most IRFs, NFkB, and CTCF. As expected, most TFs capable of initiating enhancer-wide accessibility changes at caQTLs are also predicted to drive overall accessibility (92%). The majority of them (44 or 71%) also have high context scores, indicating that binding sites for this subset of TFs are generally found in caQTL enhancers. Therefore, we coin this subset context-initiator TFs. The remaining 18 (or 29%) of initiator TFs likely reflect context-independent binding of particularly potent TFs, hence the name ‘independent-initiator’. A good example is CTCF, whose binding sites are not generally enriched in caQTL enhancers. However, when a SNP does create a new binding site for CTCF, it inevitably leads to large accessibility changes. Interestingly, we also found numerous TFs that relate to overall enhancer accessibility in LCLs, but which are not associated with caQTL SNPs, nor their sequence contexts (**Figure 1e**, red shading). These TFs overlap significantly (p.value = 1.43*10^−10^, Fisher’s exact test) with the previously identified universal stripe TFs^46^ (USTs) that provide accessibility for other TFs in a cell type-agnostic manner (**Supplemental Figure 1f**). For simplicity, we collectively refer to this class of TFs as USTs. Finally, we identified a class of TFs, including the MADS-box MEF2 and several homeodomain motifs, that lack the ability to initiate but whose binding sites are nonetheless enriched in the context of caQTL enhancers. The vast majority of them (58 or 85%) is not independently associated with enhancer accessibility, suggesting that they require context-initiator TFs for binding site access and contribute to caQTL contexts through independent, currently unknown mechanisms. We thus name them ‘context-only’ TFs.

Since condensate-like environments have been linked to cell identity genes^30,47^, we wanted to know if the two context-specific TF classes would display a certain degree of cell type specificity. To do so, we split the random enhancers into those with above or below average binding sites for a given TF and used the annotated regulatory index of DNAse I hypersensitive sites^48^ to assign enhancer annotations. Next, we assessed whether the ‘lymphoid’ to ‘tissue-invariant’ annotation ratio was larger in above versus below average binding site enhancers (**Figure 1f**). Indeed, context-initiator and context-only TFs significantly favored lymphoid-specific enhancers, whereas USTs and independent initiator TFs favored tissue-invariant elements (**Figure 1f**).

### Context-only TFs are not associated with transcriptional activity

To further investigate the functions of the four TF classes, and especially the novel class of context-only TFs, we assessed their respective contributions to transcriptional activity. For this, we chose over a thousand caQTL enhancers **(see Methods**) and measured autonomous transcriptional activity for both the more and the less accessible genotype using the SuRE-seq assay^49^ (**Figure 2a**). We found that transcriptional activity varies about 8-fold across all tested fragments, with a good overall replicate agreement (rho = 0.92) (**Figure 2b**).

**Figure 2.**
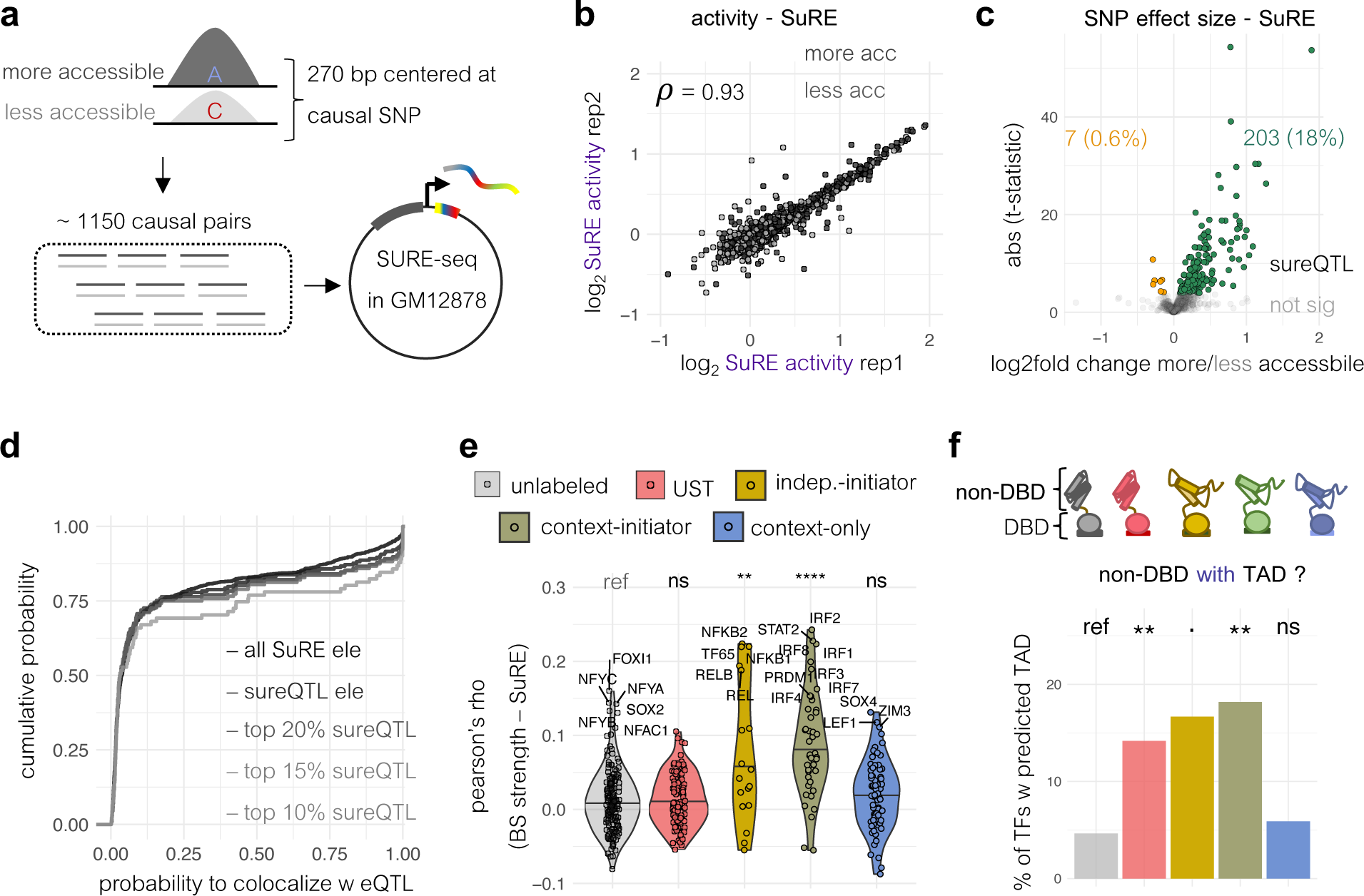
Context-only TFs are not linked to autonomous transcription. **a** Experimental overview: >1000 caQTL enhancer fragments (270bp centered around SNP) were cloned upstream of a TSS using the SuRE technology. Both more and less accessible genotypes were included. **b** Replicate agreement of average log2 SuRE activities independent of fragment orientation. Grey color scale denotes less and more accessible genotypes. **c** Volcano plot of SNP effect sizes on SuRE activity (Student’s t-test comparing less and more accessible genotypes). The Y-axis reflects the absolute t-statistic for each SNP and the x-axis the log2 fold-change in activity. SNPs with significant changes (sureQTLs) are colored (green for concordant and orange for opposite effect size direction). **d** Empirical cumulative distribution of the probability with which a SNP colocalizes with an eQTL. Gray lines represent different subsets of SNPs stratified by sureQTL effect sizes. Black line reflects the baseline of all tested caQTL fragments. **e** Correlation of fragment binding site (BS) strength (top score) with SuRE activity for each TF. Groupings reflect the four TF classes identified in Fig. 1e. Significance is computed with respect to unlabeled TFs (grey, Mann Whitney test). **f** Percentage of TFs with a predicted trans-activation domain (TAD) in their non-DBD with TFs split by TF class. Significance is computed with respect to unlabeled TFs (Mann Whitney test).

We identified 210 SNPs with differential SuRE activity (sureQTLs), or roughly 20% of all tested caQTL SNPs (**Figure 1c**). sureQTLs were more likely to colocalize with eQTLs (**Figure 2d**) and the effect size for this subset significantly correlated with the fold change in allele-specific accessibility (ASA) in GM12878 cells (**Supplemental Figure 2a**), suggesting that about a fifth of all caQTLs are driven by mechanisms related to transcriptional activity.

To investigate whether a TF may drive transcriptional activity, we compared how often a sureQTL versus a caQTL-only SNP creates a new best binding site for a TF. The 50 TFs with the highest odds ratios included several known transcriptional activators, including KLF4^50^, IRF and the NFkB/REL family. However, we detected no discernable pattern among the four TF classes (**Supplemental Figure 2b**). To more generally test a TFs contribution to transcriptional activity, we computed the correlation between fragment binding site strength (top scoring, **see Methods**) and SuRE activity independent of sureQTL status (**Figure 2e**). Both initiator TF classes, but not the context-only one were significantly correlated with SuRE activity (**Figure 2e**). To test if the observed trend would translate in actual differences in TF protein domain composition, we predicted transcriptional activation domains^51^ for non-DBD amino acids across all TFs. Compared to unlabeled TFs, we found that the two initiator classes had the highest fraction of predicted activation domains, followed by USTs. However, no enrichment was seen for context-only TFs (**Figure 2f,** Fisher’s exact test).

### Context-only TFs promiscuously pair with context-initiator TFs

Context-only TFs may contribute to caQTL environments by enhancing the function of context-initiator TFs through canonical TF complex formation. To test this, we compared the average binding site strength of context-only TFs in caQTL enhancers partitioned by whether a SNP created a new best site for a specific context-initiator TF (**Figure 3a**). Using hierarchical clustering (**Figure 3b**), we indeed detected a cluster of context-only and context-initiator TFs that belong to a previously described, cooperatively acting pair – FOXO (context-only) and Ets (context-initiator) TFs^9,52^ (**Figure 3b**, orange cluster). FOXO sites, however, are not specific to Ets factors, since their binding sites also score high in caQTL enhancers with newly gained RUNX sites. In general, the majority of context-only TFs paired with multiple context-initiators. This was particularly true when the pair included non-Ets context-initiators (**Figure 3b,** blue clusters). Importantly, we did not find a similarly pattern when replacing either context-initiator or context-only TFs with USTs (**Supplemental Figure 3a&b)**. This indicates that the pairing of context-only and context-initiator TFs is not exclusive to specific motif combinations, but applies more loosely to combinations of TFs with complementary function, a concept in line with ‘soft’ motif syntax rules^12^.

**Figure 3.**
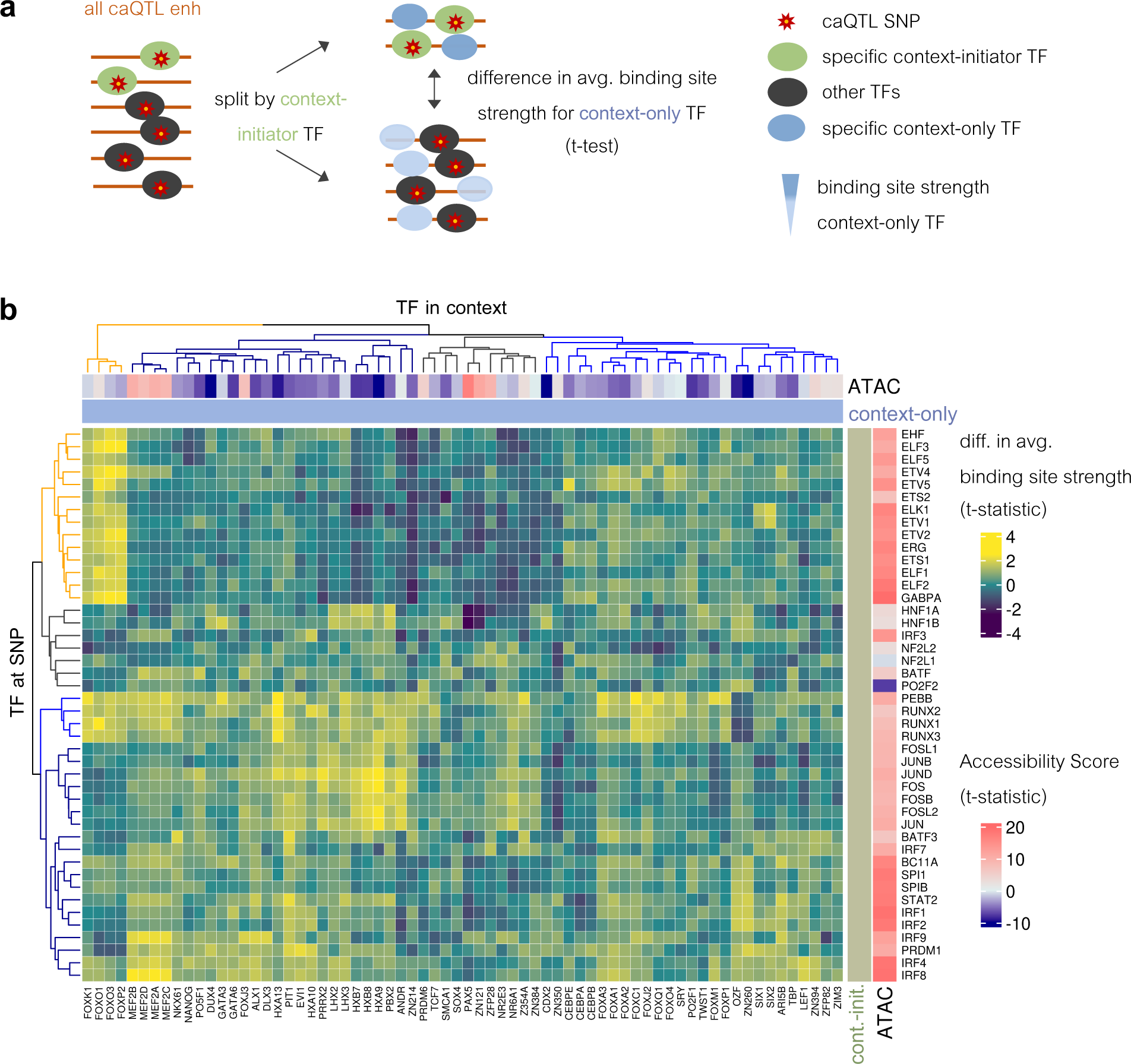
Context-only TFs pair promiscuously with context-initiators. **a** Schematic describing how combinatorial TF binding is assessed: for each context-initiator TF caQTL enhancers are split based on whether the SNP creates a new best site or not. Next and for each context-only TF separately, the binding site score distributions are compared between caQTL enhancers split by context-initiator binding site strength (Student’s t.test). **b** Hierarchical clustering based on the t-statistic computed for each context-initiator/context-only pair described in **a** and split in four main clusters (dendrogram colors). Columns contain context-only and rows context-initiator TFs. The orange cluster contains previously identified cooperative pairs. A TF’s accessibility score is shown in the outermost annotation panel (blue-red color scale).

### Context-only TFs induce cooperativity in enhancer activity assays

Given their promiscuous pairing, we next hypothesized that context-only TFs may induce cooperativity in a promiscuous manner. To experimentally test this, we chose 8 representative motifs, two context-only (MEF and FOX), three context-initiator (SPI, IRF, RUNX), two independent-initiator (NFkB, CTCF), and one unlabeled motif (MYC) and tested how homo- and heterotypic motif combinations affect intrinsic enhancer activity using the STARR-seq assay^53^ (**Figure 4a**). To account for positioning effects and motif-flanking sequence, we included two different spacers (5bp and 10bp) and embedded each motif combination into three randomly created and barcoded sequence contexts (**Figure 4a** & **Supplementary Table 1**). We measured enhancer activity for a variety of motif combinations, including up to five homotypic repeats, all pairwise combinations, and all 3-motif combinations of two TFs (**Figure 4a**). We found that motif identity, spacer length, and motif number all influence enhancer activity (**Figure 4b and Supplemental Figure 4c**). For several TFs, particularly in the 10bp library, we found that enhancer activity trended down with each motif added (**Figure 4b** and **Supplemental Figure 4c**). Focusing on cooperativity, we assessed whether adding a context-only to a context-initiator motif would lead to an increase in activity compared to each motif on its own or in combination with non-context-only motifs. As an example, we took MEF2 and PU1, and compared enhancer activity across all homotypic or heterotypic pairings of these two motifs (**Figure 4c**). Strikingly, the combination of MEF2 and PU1 motifs resulted in enhanced activity compared to each motif on its own (**Figure 4c**). We found similar enhancing effects for RUNX and FOX motifs (**Supplemental Figure 4a**). Intriguingly, the combination of the two context-only motifs (FOX and MEF2) resulted in even stronger increases in enhancer activity, suggesting that their combined binding leads to productive cooperativity in the context of episomal DNA (**Figure 4c).** Importantly, no effect was found when combining different context-initiator TFs (**Supplemental Figure 4b**), supporting their primary role in providing DNA access, or mediating transcriptional activity, but not cooperativity per se.

**Figure 4.**
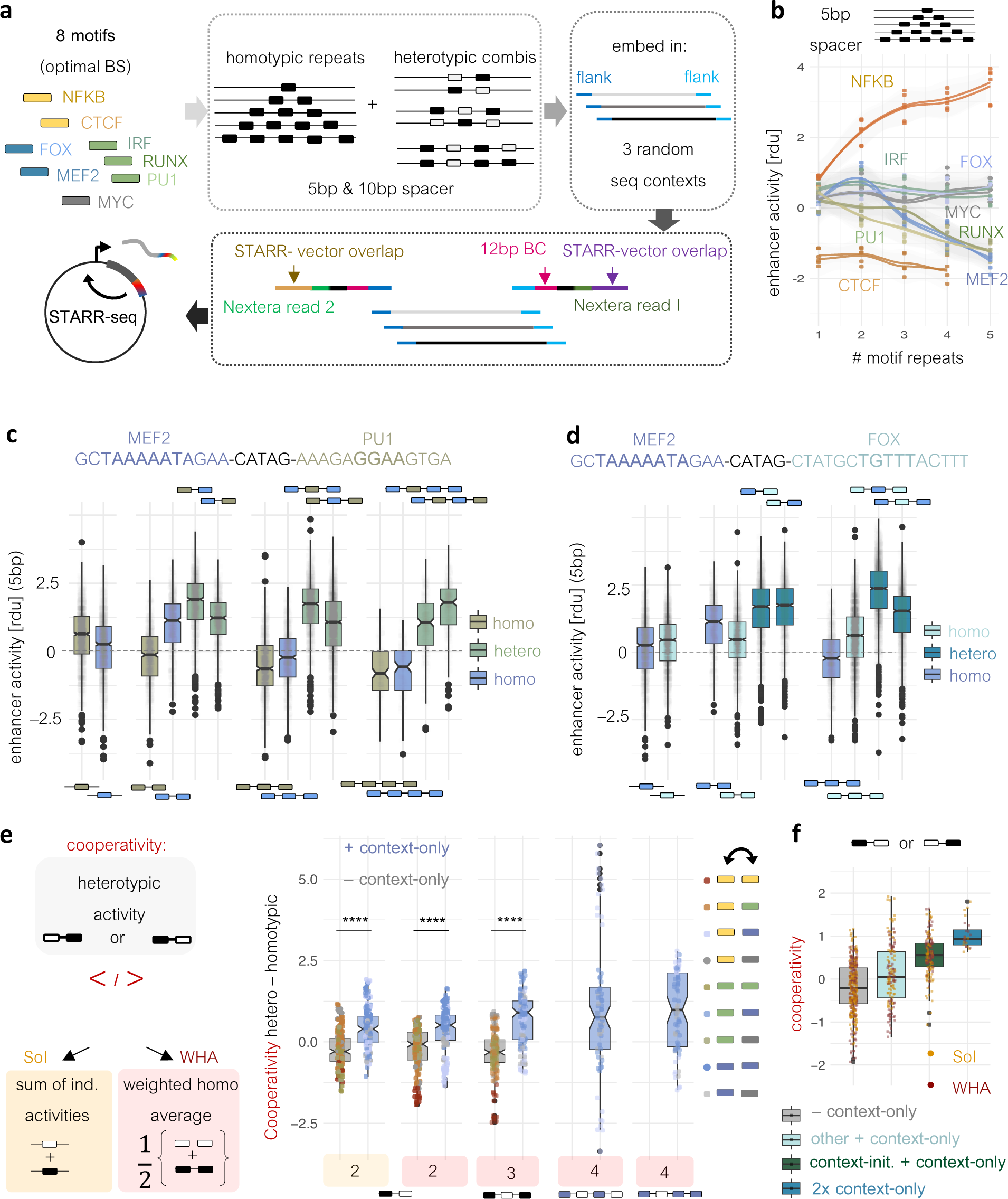
Context-only TFs induce cooperativity in enhancer activity assays. **a** Schematic description of the experimental setup: 8 different motifs were embedded in 3 different, random sequence contexts in form of homotypic and heterotypic motif repeats with two different spacers. Each fragment was tagged by PCR with a 12bp random barcode and cloned into the STARR-seq vector. **b** STARR-activity for homotypic motif repeats measured as the average across all barcodes and normalized using the respective motif-free random sequence as a reference (rdu = in units of random context). Colors reflect the TF motif classes defined in Fig. 1e. Lines are separate replicates and points belong to one of the three random sequence contexts. **c** STARR activity for individual barcodes of homotypic or heterotypic motif combinations of PU1 and MEF2 normalized by the average STARR activity of the respective random sequence contexts (y-axis). Motif orientation and 5bp-spacer sequence is given on top. The exact motif composition is indicated below (homotypic repeats) or above (heterotypic repeats). Colors indicate the TF (blue = MEF2; yellow green = PU1, blue-green = MEF2 & PU1). **d** Same as **c** but for the combination of MEF2 and FOX motifs. **e** Schematic describing how cooperativity is computed (left): Sum of individual (SoI, orange) compares the average activity of a heterotypic motif combination versus the sum of the individual motifs. WHA (red) compares against the weighted average of homotypic motifs with the same number of motifs. Right: Cooperativity (y-axis) in the 5bp spacer library as a function of motif number (x-axis schematic), colored by the method (SoI or WHA) below each individual comparison. Color of boxplots indicate whether a motif combination contains a context-only TF or not (blue/grey). Color of points indicate different combinations of TF classes (class color corresponds to the motif color in **a**. Different sequence contexts and replicates represent individual points. **f** Same as **e** but focusing only on two motif combinations and splitting based on distinct TF class combinations (x-axis). Boxplot colors indicate class combinations. Points are colored based on the methods used to assess cooperativity described in **e.**

To systematically assess the cooperative effect provided by context-only motifs, we derived two measures of cooperativity, comparing heterotypic enhancer activity either to the sum of individual motifs or the weighted sum of homotypic motif combinations of equal length (**Figure 4d).** Heterotypic motif combinations that included context-only motifs were associated with positive cooperativity and scored significantly higher than those without (**Figure 4e**). This was true independent of motif number and spacing (**Figure 4e and Supplemental Figure 4d).** Splitting heterotypic motif pairs that contained a context-only motif based on the class of the second motif, we found that the two context-only motifs do particularly well when paired with context-initiator motifs or each other (**Figure 4f and Supplemental Figure 4e)**.

### Context-only TFs have low complexity domains and enhance BRD4 recruitment

Given their ability to induce cooperativity, we hypothesized that context-only TFs contribute to the creation of caQTL environments by tethering other TFs and potentially coactivators to enhancers targeted by context-initiator TFs, thus creating localized RF clusters. In this case, context-only TFs may engage in weak protein interactions mediated by disordered domains. To test this, we compared the fraction of non-DBD amino acids with low complexity^54^ (a proxy for disorder) across the four motif classes (**Figure 5a**) and indeed found a higher fraction in context-only TFs compared to USTs, context-initiators, and even the bulk of unlabeled TFs (**Figure 5b**).

**Figure 5.**
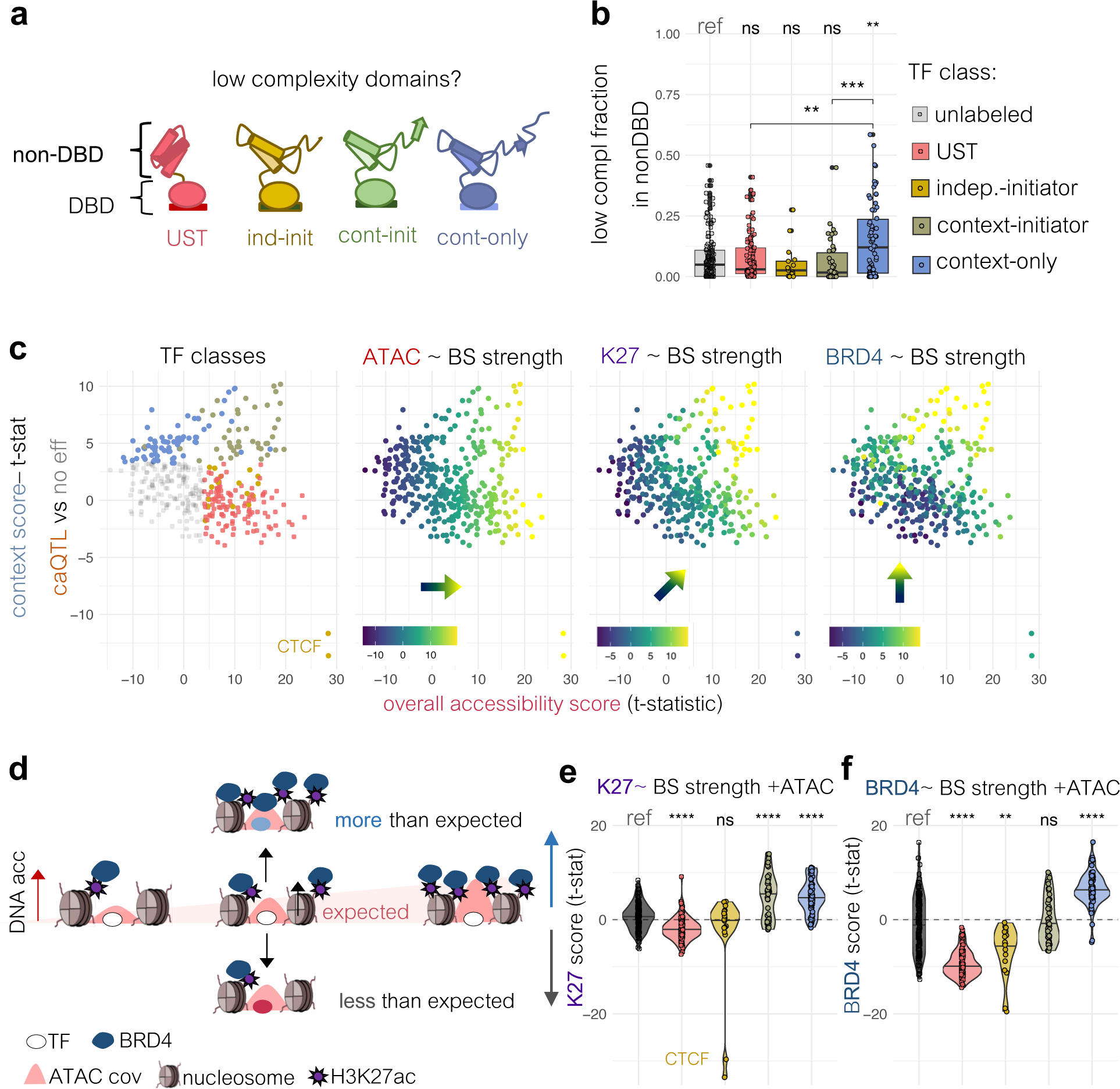
Context-only TFs are enriched for low complexity domains and recruit BRD4. **a & b** Fraction of 20-aa windows with low complexity (y-axis, B) within the non-DBD of TFs (schematic in **a**) across TF classes. Significance is computed using unlabeled TFs as a reference (Mann Whitney test). **c** TF context score (y-axis; cf. Fig. 1d) versus TF accessibility score (x-axis; cf. Fig. 1d) colored in order by the TF class, the raw accessibility score (same as x-axis), and the binding site (BS) strength association of each TF with either H3K27ac or BRD4 ChIP-seq coverage in GM12878 (t-statistic of linear model coefficient). Blue-yellow color scale represents negative and positive associations. **d** Schematic explaining the model used in **e/f**. Accessibility drives baseline levels of H3K27ac and BRD4 IP coverage (red shaded area represents the expected enrichment). TFs are evaluated on whether their binding site strength (top score) is associated with more or less IP coverage after controlling for accessibility. **e & f** Results of log-linear model associating IP coverage (H3K27ac = **e**; BRD4 = **f)** with binding site strength while using ATAC-seq coverage as a covariate. T-statistics of the corresponding coefficients are plotted on the y-axis. TFs are split by TF classes and significance is assessed with respect to unlabeled TFs (Mann Whitney Test).

Next, we wondered how context-only TFs relate to the recruitment of coactivators. We focused on BRD4, as it harbors a strong IDR^26^, is linked to condensate formation, and can recognize acetyl lysine residues on H3K27 through its bromodomain. Specifically, we used the randomly sampled LCL enhancers and tested how TF binding site strength relates to either the overall ATAC-seq, H3K27ac- or the BRD4-ChIP-seq coverage (log-linear model, **see Methods**). Overlaying the coefficients of each TF-to-epigenomic state model onto the space spanned by the context and accessibility scores (cf. Figure 1), we uncovered a distinct pattern: while accessibility is strongest for both USTs and context-initiator TFs (**Figure 5c**), H3K27ac- and BRD4-enrichment shifts away from USTs towards context TFs, with the TF-BRD4 association gradient almost perfectly aligning with the “context score” axis (**Figure 5c**). Since DNA accessibility correlates with many molecular marks, including H3K27ac and BRD4, we recomputed the relationships while including accessibility as a separate covariate (**Figure 5d**). Doing so, we found that UST binding strength is associated with lower, while that of context-only and context-initiator TFs with higher H3K27ac levels (**Figure 5e**). Intriguingly, the context-only class remained the only class significantly associated with BRD4 recruitment (**Figure 5f**).

### Enhancer communication is linked to context-TF enrichment

If context-only and context-initiator TFs indeed cooperate to locally enrich BRD4 and create RF clusters, their binding to two or more enhancers may result in fusion of the two clusters into a shared hub^39,55^. To test this hypothesis, we used information on CMs provided in the original caQTL study^43^. Briefly, the study applied a Bayesian hierarchical approach^43^ to infer CMs and their internal hierarchies, such as when the accessibility change of a caQTL ‘lead’ peak is inherited by at least one ‘dependent’ peak (**Figure 6a**). To infer which TFs contribute to such enhancer coordination, we compared lead and dependent enhancers to a control set of independent enhancer pairs, which consist of a caQTL enhancer (LOCAL) and its unlinked neighbor (**Figure 6a and Supplemental Figure 5a**). To minimize confounders, we matched caQTL probabilities, as well as GC content between LOCALs and CM-leads using fifty separate samplings (**Supplemental Figure 5a&b).**

**Figure 6.**
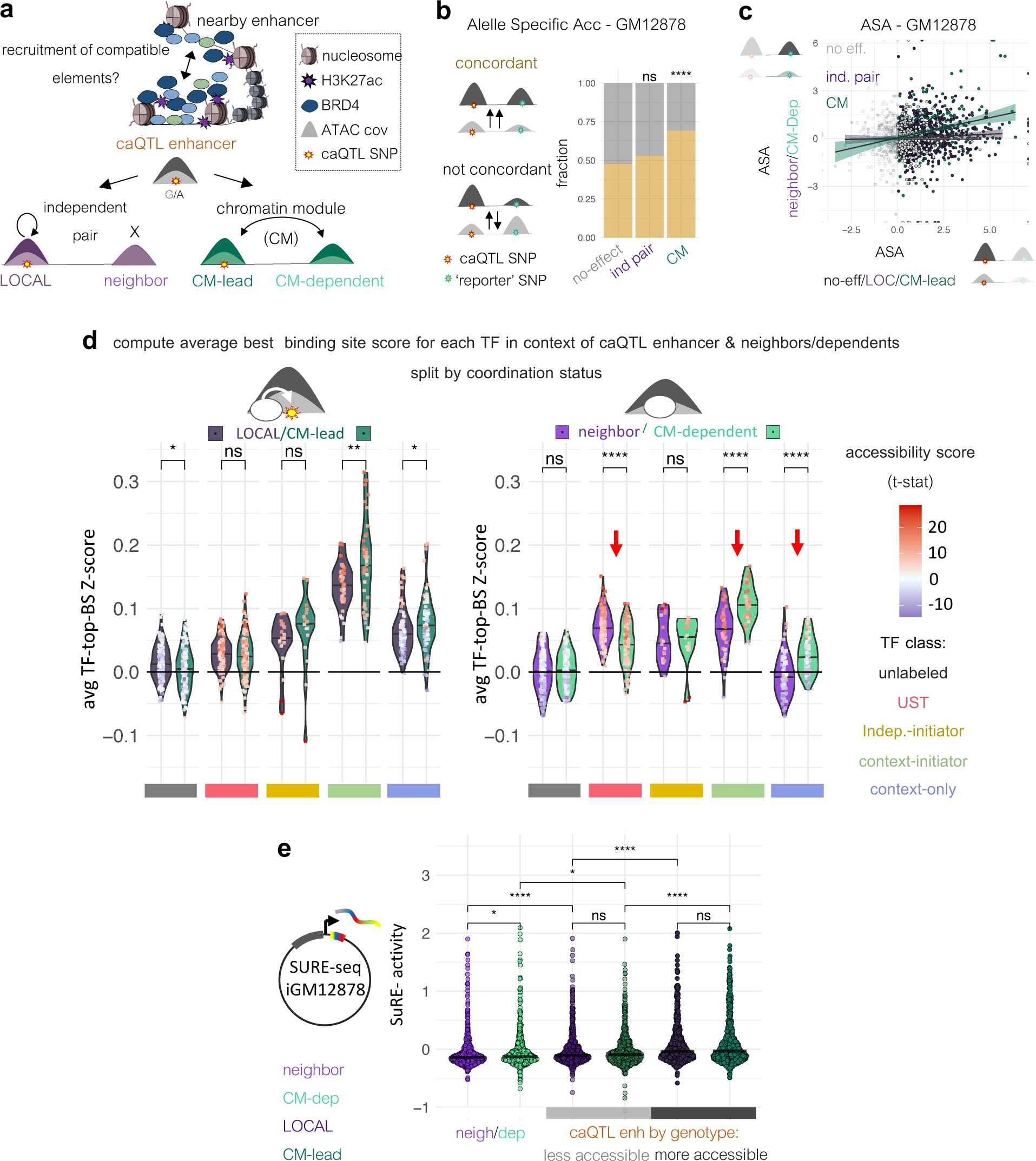
Context-TFs explain CM formation. **a** Schematic illustrating CM formation based on context-TFs and BRD4 (top) and how they are discovered using DNA accessibility readout at the population level (bottom). caQTL enhancers fall into two groups: Those with neighboring enhancers that do not display accessibility changes associated with the caQTL genotype (independent pairs, purple), and those whose accessibility change is inherited by a nearby element as a function of caQTL genotype (CM-lead and CM-dependent, sea green). Less and more accessible genotypes are indicated by dark and light color shading. **b** Validation of independent pairs and CMs in GM12878 using phased genome data. Concordance of allele-specific accessibility changes (ASA) is assessed by comparing the direction of change at heterozygous caQTL (red star) and reporter SNPs (blue star, in neighbors or dependent enhancers) in GM12878. Concordant/non-concordant changes in yellow/grey. Significance is assessed for independent pairs and CMs by comparing the respective fractions to no-effect enhancers and their neighbors (Fisher’s exact test). **c** Relating the effect size of ASA at the reporter SNP (**b**) to the ASA at the caQTL SNP as a function of enhancer type (no-effect = grey; independent pairs = purple, CMs = sea green). **d** Average TF binding site scores (y-axis; top scores excluding SNP, see schematic on top) expressed as Z-scores using the score distribution of randomly sampled enhancers as a background. Average Z-scores are split by enhancer sub-categories and TF class. LOCAL and CM-lead caQTL enhancers are shown on the left with darker shadings and neighbors/dependents on the right with lighter shading (LOCALs/neighbors = purple; CM-leads/-dependents = sea green). TF classes are indicated as colored boxes (x-axis). Black lines represent the expected average Z score in the random enhancer sample. Significance is assessed by comparing within enhancer categories and within a given TF class (Mann Whitney test). Red arrows indicate significant class-wide differences in average binding site scores in neighbor/CM-dependent enhancers. **e** SuRE-seq data for independent and CM enhancer pairs, y-axis indicates the log2 SuRE activity averaged across all barcodes and orientations for an individual fragment. Shading differentiates caQTL from neighbors/dependents and the color differentiates independent from CM pairs. For caQTL enhancers, the two different genotypes (less and more accessible) are indicated by differently shaded grey bars on the x-axis. Significance is assessed across enhancer categories (Mann Whitney test).

First, we validated that CMs exist in GM12878 cells, where trans effects, e.g., differences in TF expression levels across individuals, can be excluded. Using phased genome information, we computed the concordance in allele-specific accessibility (ASA) at the CM-lead or LOCAL SNP with that of a reporter SNP within dependent or independent neighbors. We found that only SNPs linked through a CM show a significant concordance (**Figure 6b**) with a roughly linear relationship with ASA effect size (**Figure 6c**). The latter was true, even when including enhancer distance, reporter SNP to peak center distance, as well as label free ASA at the caQTL enhancer as covariates in a linear model (**Supplemental Figure 5c**). Next, we ruled out that differences in chromatin topology may account for CM formation by comparing contact frequencies between distance-matched independent and CM enhancer pairs, finding no significant difference (**Supplemental Figure 5d&e**).

Finally, we assessed whether CMs differ in their overall binding site composition. While no differences were found when comparing the frequency with which SNPs create new best sites (**Supplemental Figure 5f**), LOCALs and CM-leads and their respective neighbors and dependents differed in their average context binding site scores across TF classes. We found that on average both context-initiator and context-only TFs have significantly higher binding site scores in CMs compared to independent pairs (**Figure 6d**), with the differences more pronounced and more significant when comparing LOCAL-neighbors with CM-dependents. It also revealed a reverse selectivity for USTs, whose binding sites were stronger in independent neighbors (**Figure 6d)**. Context-only TFs are of particular interest, as their average binding site strength surpassed that of the random enhancer sample only in CM-dependents but not in LOCAL neighbors (**Figure 6d**).

Interestingly, the hierarchy within CMs based on Bayesian modeling of ATAC-seq coverage is recapitulated in the average motif sores for context-associated TFs, with lead enhancers having higher averages compared to dependents (**Figure 6d**). To assess whether this binding site hierarchy also translates into activity differences, we analyzed the SuRE-seq data split by caQTL enhancer status (CM-lead or LOCAL). In addition, we included the corresponding neighbor/dependent enhancers into the assay. Median activities varied across groups, with lower medians observed for neighbors and dependents compared to their respective LOCALs and CM-leads. This was true even when comparing to the less accessible caQTL genotype, suggesting that hierarchies are innate and not gained *de novo* by the introduction of a variant **(Figure 6e**). There was no difference when comparing LOCALs and CM-leads. However, CM-dependents scored higher than LOCAL-neighbors, perhaps reflecting the higher average binding site scores for context-initiator TFs which we could link to autonomous transcription (**cf. Figure 6d & Figure 2e**).

### Genes controlled by CMs are sensitive to BET inhibition

Since enhancers linked through CMs resemble SEs, they should be similarly sensitive to Bromodomain and Extra-Terminal domain inhibitors (BETis)^56–58^, i.e. JQ1 (**Figure 7a**). BETis not only disrupt gene expression controlled by SEs, but also do so without changing the overall contact frequency between REs^59^, suggesting that they specifically target RF clusters, which we hypothesize includes those created by context TFs.

**Figure 7.**
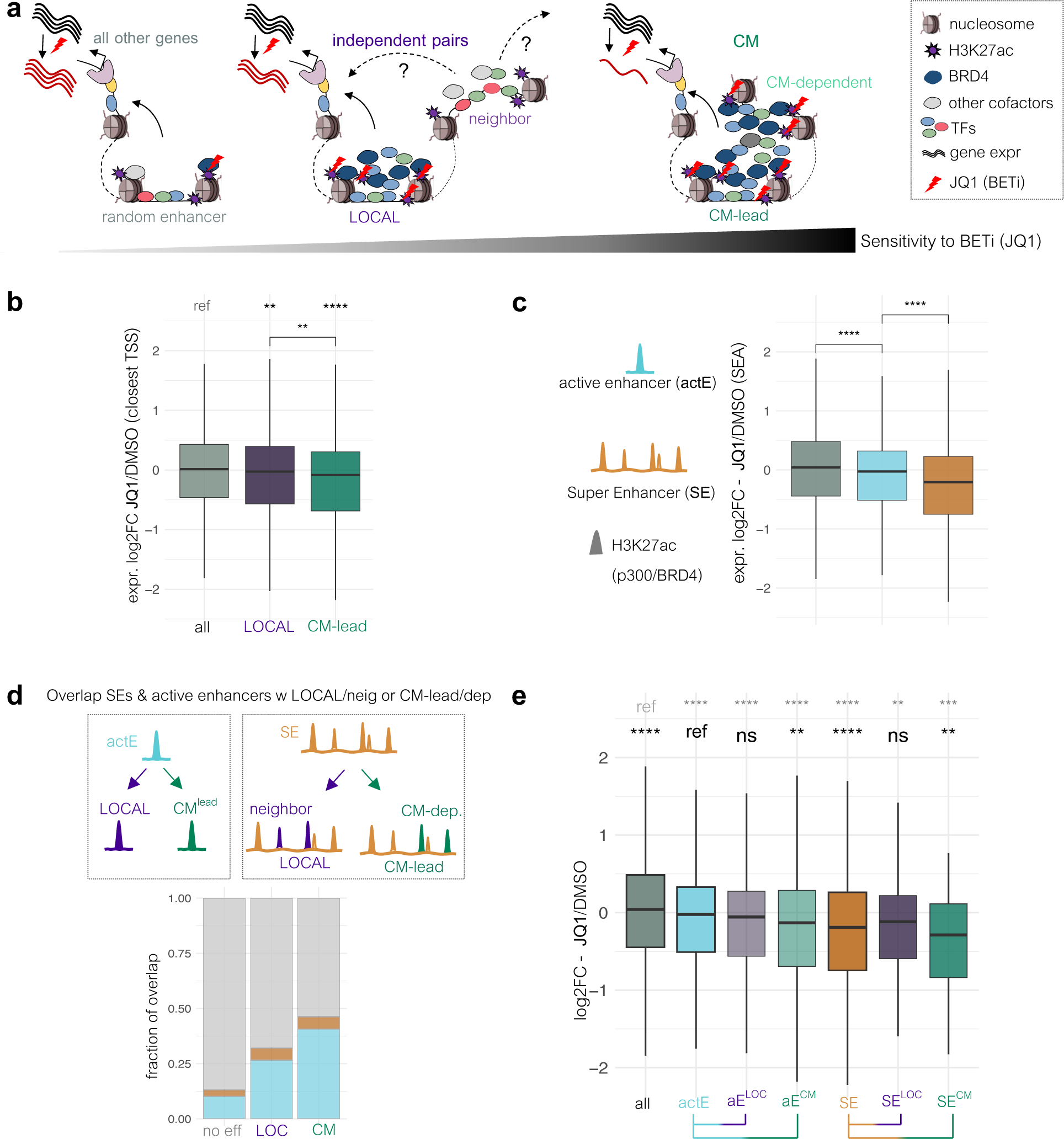
Genes controlled by CMs are sensitive to BET inhibitor JQ1. **a** Illustration of the impact on gene expression exerted by different molecular assemblies at enhancers: random enhancers with unknown regulatory status (left), LOCALs with independent neighbors (middle) and CMs (right). From left to right, the combination of context-initiator and context-only TFs leads to more H3K27ac and BRD4 enrichment and thus heightened sensitivity to JQ1 (color bar below). JQ1 disruption of the bond between BRD4 bromodomains and acetyl lysine residues at H3K27, as well as the expected effect on gene expression is indicated by red lightning signs. **b** Log2-fold-change (log2FC) in expression after JQ1 treatment of MEC1 cells, comparing genes controlled by LOCAL and CM-lead caQTL enhancers to the baseline of all other genes. Enhancer are mapped to their closest gene promoter. **c** Same as in **b** but using the enhancer and gene annotations provided in SEA. **d** Overlap between caQTL or no-effect enhancers with active B-cell enhancers (actEs; cyan) and independent/CM pairs and no-effect-neighbor pairs with SEs (orange). Non-overlapping enhancers in grey. **e** Same as in **b/c** but splitting active enhancers based on their LOCAL/CM-lead annotation and SEs based on their independent- or CM-pair label. Significance is assessed with respect to the baseline of all (grey) or active enhancer genes (black) using a Mann Whitney test.

Using existing gene expression data of LCLs treated with either DMSO or JQ1^56^, we asked whether genes controlled by LOCALs or CM-leads display sensitivity to JQ1 and if yes, to what extent. Despite some caQTL enhancers likely not controlling gene expression directly and using a simple ‘closest TSS’ enhancer-to-gene mapping approach, we nonetheless found that both LOCAL and CM-lead genes are sensitive to JQ1, with the latter having a larger impact (**Figure 7b**). There was no difference in expression levels among the two groups (**Supplemental Figure 6a**). To compare CMs directly to SEs, we first acquired recently updated enhancer annotations from the Super Enhancer Archive^60^, including both active B-cell enhancers and SEs. The latest annotations revealed that active enhancers are also sensitive to JQ1, albeit to a lesser extent than SEs (**Figure 7d**), thus resembling the sensitivity gradient of LOCAs and CM-leads. Baseline expression for genes in both active and SEs was skewed towards high expression (**Supplemental Figure 6b**), which likely reflects the discovery process that relies on high levels of active marks^29^. Enhancer communication, however, may not require constitutively high activity and vice versa, high activity alone may not imply communication. For instance, independent enhancer pairs may still classify as SEs, despite communicating separately with potential target genes. (**cf. Figure 6d & 7a**). To investigate the sensitivity of high H3K27ac-versus CM-derived annotations, we first identified the overlap between CM and independent pairs with either active enhancers or SEs (**Figure 7d**). We found that compared to LOCALs, CMs are enriched for active enhancers (**Figure 7d**), suggesting that the presence of a dependent enhancer may increases the deposition of active marks (**Figure 7a**). Importantly, we did not detect differences in SE overlap, which indicates that the current SE label may not sufficiently differentiate between true coordination and independent activity. Indeed, when comparing JQ1 sensitivity among the different subcategories, we found that only the CM-, but not the LOCAL-label displays a significantly enhanced sensitivity when comparing to active enhancer-genes (**Figure 7e**). Moreover, CM-active enhancer and CM-SE genes had larger median reductions in gene expression upon JQ1 treatment than the LOCAL, as well as the remaining active and SE genes (**Figure 7e** & **Supplemental Figure 6c**). This suggests that the signature for context-TFs, in particular when present in two or more CM-participating enhancers, provides a mechanistic framework for the sensitivity to JQ1which adds granularity to the classical SE label.

## Discussion

RF clusters, transcriptional condensates, Super Enhancers, or phase-separated hubs are all terms that describe nuclear environments with elevated RF concentrations. Here we describe a new approach that allowed us to shed light on the TF-mediated mechanisms that underlie the formation of such clusters. By considering the non-linear recruitment process of molecules into growing condensates^35,38,39^ and equating it to the phenomena observed at enhancer-centric caQTL SNPs, we were able to identify a novel class of TFs, context-only TFs, whose function directly relates to the creation of cooperative environments.

Context-only TFs are fundamentally different from initiator TFs that are found at causal variants themselves. Their binding site strength is not generally associated with DNA accessibility, arguing that they lack any pioneering ability. Rather, context-only TFs are found in the vicinity of a subset of initiator TFs with cell type-specific binding signatures. In addition, context-only TFs contain low complexity domains, induce cooperativity with flexible motif pairing and syntax, and their binding site strength is associated with above-average BRD4 deposition. All these findings are in line with a model by which the creation of a high RF, condensate-like environment, rather than independent binding of extremely potent TFs, leads to the majority of large effect sizes observed at enhancer-centric caQTLs. As such, our findings provide a mechanistic rationale for the action of pioneer and ‘lineage’-specifying TFs^61^: cell type specificity relies on the combined action of two types of context-specific TFs. One has a pioneering role and the other an amplifying one. Such a mechanism may explain non-canonical pioneers, such as PU1^62^, that rely on genomic context to provide access to DNA. Indeed, PU1 is among the TFs labeled as context-initiators whose activity at SNPs is linked to context-only TFs such as MEF2.

That this dual function mechanism is neither restricted to caQTLs, nor B-cells, but rather applies more generally is supported by two recent studies. In one study, *D. melanogaster* patterning TFs enhanced accessibility and marked developmentally active enhancers when combined with the pioneer Zelda^63^. Although the exact molecular mechanism remained elusive, the role of patterning TFs closely matches the one of context-only TFs. Another study in yeast showed that non-DBD TF regions, particularly those in disordered domains contribute to TF target specificity^64^, providing evidence that the low complexity domains in context-only TFs may indeed license TF localization. Nonetheless, studies in additional cell types will be required to test how generalizable the context-TF driven mechanisms are, and whether context-only TFs are true lineage-specifiers or perhaps simply adapt to new, cell-type specific initiators.

For now, we can only speculate how exactly context-only TFs establish cooperativity. The simplest explanation would be nucleosome-mediated cooperativity, by which multiple TF binding sites act allosterically to evict nucleosomes^65^. However, several of our findings indicate that such a model does not suffice: For once, we might suspect USTs that were previously shown to grant access to DNA for other TFs^46^ to adhere to the same mechanism. Yet, their binding sites are not found at cis-acting causal variants. More importantly, nucleosome-mediated cooperativity does not require a link to the histone code nor an enrichment for disordered non-DBDs. It neither explains enhancer communication, nor the enhanced sensitivity to BETis. The creation of condensate-like RF clusters, however, fits all criteria.

A plausible order of events could look like the following: Context-initiators initiate the removal of an additional nucleosome, which is then stabilized by the binding of context-only TFs. The critical component however is the establishment of H3K27ac marks, followed by the binding of BRD4 to acetyl lysine residues. BRD4 recruitment is then stabilized by context-only TFs and likely involves their disordered non-DBDs. When two enhancers with similar context-TF driven environments are in proximity, the two RF clusters fuse, resulting in a transient communication^66^ that amplifies effects on downstream gene expression, thus rendering genes sensitive to BETis. Such a scenario would rely on the cumulative action of TF DBDs and non-DBDs without the requirement for distinct combinations and exact motif syntax and would rationalize why cracking the cis-regulatory code remains a challenge.

Finally, context-TF binding explains enhancer compatibility at a molecular level, a conceptual task that has remained challenging in the past^67^. Our finding that enhancer independence is associated with the class of USTs provides further evidence that individual enhancers can achieve DNA accessibility through fundamentally different mechanisms. Importantly, our findings suggest that the decision to communicate lies predominantly with the ‘weaker’ (dependent) element.

Dependent enhancers thus resemble the recently discovered ‘facilitator’ elements that are part of the erythroid a-globin SE^68^, as well as the ‘tethering’ elements found in Drosophila^69^. We propose that CM-dependents and facilitators or tethering elements are two sides of the same coin: they all tune the activity of another coordinated element. We thereby postulate that the underlying mechanism directly relates to the ability of TFs to create environments with high RF concentration.

In summary, we describe a new approach that leverages specific caQTLs to identify the TF-driven mechanisms underlying RF cluster formation. Abstracting to average effects allowed us to identify a global mechanism that up to this point, had remained hidden. The downside of this approach is that it does not allow for predictions at the level of individual enhancers, nor does it provide granularity on the types of cooperative environments. It is likely that future studies will reveal distinct codes for selective partitioning, similar to what has been observed for coregulators such as BRD4 and MED^70^. We thus expect that the unifying mechanism presented here will serve as a steppingstone for future experiments and lead to the development of machine learning models that can accurately predict cooperative environments.

## Code and Data availability

Raw and processed sequencing data for STARR-seq and SuRE-seq experiments, as well as processed summary statistics for enhancer pairs, and TF classifications will be made available upon publication.

Custom code will be deposited to GitHub or is available upon request.

## Author Contribution

J.F.K and B.D. conceived and designed the study. J.F.K and J.R. performed the experiments.

J.F.K analyzed the data and performed the statistical analyses, with help from O.P., V.G., and G.V.M.. J.F.K and B.D. wrote the manuscript with input from V.G., G.V.M. and O.P.

## Acknowledgements

We thank Dr. Harmen Bussemaker and Dr. Carles Canto for reviewing the manuscript and providing valuable feedback. We further thank ANNOGEN, the EPFL gene expression core facility, as well as the EPFL’s Scientific IT and Application Support (SCITAS).

This work was supported by a Swiss National Science Foundation grant (no. 310030_197082), Marie Skłodowska-Curie fellowships for J.F.K. (no. 895426), O.P. (no. 860002), and G.v.M. (no. 101026623), as well as EMBO long-term fellowships for J.F.K. (1139-2019) and G.v.M. (2020-895).

## Methods

### caQTL data pre-processing

Information on caQTLs, peaks, and CMs (called ‘DAGs’ in the original publication) in LCLs from 100 individuals were retrieved from Zenodo linked to the following publication (Kumasaka et al.^43^ DOI: 10.5281/zenodo.1405945; ‘all.caQTL.summmary.statistics’, ‘peaks’, and the ‘dag’ files). First, we merged the caQTL.summary and the peak files, keeping each caQTL instance separately. To single out SNPs that fall within the peak boundaries, we required the ‘Inside_Peak’ column to be 1 and the genotype indicated in ‘Ref’ and ‘Alt’ to be exactly 1 nucleotide in length.

### Nominating caQTL, no-effect and random enhancers

To focus on SNPs with large effect sizes, we first applied a filter requiring the following criteria: P_caQTL*P_Lead>0.7 and P_caQTL>0.999. P_caQTL is the probability for a SNP to affect the accessibility of a given peak and ‘P_Lead’ the probability of the SNP being the lead variant. We considered enhancers above this cutoff to be high confidence caQTL enhancers. We further removed all enhancers whose SNPs fell outside the +/- 350 bp around the peak center we considered for TF binding site scoring. For ‘no-effect’ enhancers, we required P_caQTL< 0.2 and P_Lead<0.5 with their product being less than 0.1. We also removed ‘no-effect’ genes that were contained within the ‘dag’ file (contains CM-annotations), as well as those that are direct neighbors of other caQTL enhancers. Next, all retained peaks were annotated with the annotatePeak function from the R package ChiPseeker^71^ using the TxDb.Hsapiens.UCSC.hg19.knownGene annotation package with 500 bp up- and downstream to indicate the TSS region. We retained all peaks that were annotated either as ‘Distal’ on ‘Intron’ to represent putative enhancers.

To avoid sequence composition biases that could skew the TF binding site inference, we downsampled the no-effect enhancers to a subset with equal GC content distribution to that of the caQTL set. We also matched the ‘SNP to peak center’ distances to avoid comparing against SNPs whose effect sizes are negligible simply because they are located at the fringes of peaks. Sampling was done by randomizing the respective values in the caQTL set and then iteratively retaining the closest match in the no-effect set. This process was repeated 50 times to generate 50 sets of controls to maximize robustness of results. Each time the order within the caQTL enhancers was randomized using a new seed, resulting in different selections of ‘no-effect’ enhancers as a result of the iterative process – the first caQTL peak in order is matched first before moving to the next. A random sample of enhancers was generated by sampling 25,000 peaks from the full peak table without replacement. Thereafter, only enhancers annotated as ‘Distal’ or ‘Intron’ were retained resulting in ∼21k total putative enhancers.

### ATAC-seq data processing and peak scoring

To identify generalizable mechanisms, we used the commonly used reference cell lines for LCLs GM12878. DNA accessibility (ATAC-seq) data for GM12878 was retrieved from ENCODE (Experiment: ENCSR637XSC). Read1 and Read2 fastq files were first combined before aligning to the hg19 reference genome (GRCh37, release 75 from Ensembl) using the bwa-mem^72^ with standard settings. Alignment files were sorted and indexed with samtools^73^ and subsequently transformed to the ‘bigwig’ format using the bamCoverage command from deepTools^74^. To obtain an accessibility value for each enhancer, all counts falling within 700 bp, +/– 350bp around the peak center (same sequence used for binding site scoring), were added up.

Human TF motifs were downloaded from the HOCOMOCO database^3^ (version.11) in forms of mono-nucleotide scoring matrices. Note, extracted raw scores from such scoring matrices reflect log-transformed relative TF affinities. To generate an enhancer context score, each 700 bp sequence was scored in forward and reverse orientation using the ‘calculate’ function of the Biopython package^75^ (Bio.motif). For each enhancer the best score was retained as the final enhancer binding site score (top scoring), as it reflects the largest attraction potential for a given TF within an enhancer. For caQTL enhancers, scoring and top score extraction was done for each genotype separately (less and more accessible based on the sign of the SNP effect size coefficient – Beta). If the two resulting top scores were different, the SNP either created a new or destroyed an existing best site. In addition to scoring actual enhancer sequences, we also scrambled the 700 bp sequences of random enhancers and recomputed the top scores in the same manner. The resulting scrambled top scores are used whenever sequence composition bias, i.e., GC content bias intrinsic to experimental methods such as ATAC-seq^76^, may confound the inference of TF binding site preferences. For example, if a scoring matrix for a TF contains a high scoring CG, it is more likely to produce a high-ranking top score in CG-rich regions even if the full motif itself is not enriched.

### Accessibility, initiator and context-score computation and thresholding

To compute the association of a TF with overall accessibility (accessibility score), we used a log-linear regression model, relating the cumulative ATAC-seq signal at a given enhancer to a TF’s top score. We used the top scores of scrambled sequences for the same TF as an additional covariate and fit the following model for each TF separately in R:

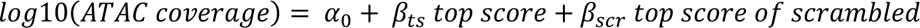

With α_3_, β_67_, β_7;<_ being the TF-specific intercept, and the coefficients relating overall accessibility of an enhancer to a TFs top score in the actual or scrambled enhancer sequence. We extracted the corresponding t-statistic of β_67_to represent a TF’s accessibility score. To define TFs with a significant association with accessibility we adjusted the p-values using the ‘Holm-Bonferroni’ correction and set the significance cutoff to p<0.01, resulting in a final t-statistic cutoff of t >= 4.

To obtain the initiator score, we first computed how many times a SNP created a new best site for a TF when going from the less to the more accessible genotype. We did this separately for caQTL and no-effect enhancers. Next, using Fisher’s exact test, we computed the odds ratio comparing the number of events in caQTL versus no-effect enhancers. We repeated this approach for all the 50 samples of ‘no-effect’ enhancers. For each TF a final log2 odds ratio was assigned by averaging the results across the 50 samples. The cutoff for the log2 odds ratio of 0.9 was set by using the scatter along the ‘more frequent in no-effect enhancer’ direction as a reference for what would be expected due to pure chance (there is no reason to believe that a randomly sampled SNP that causes minimal to no change should specifically attract a TF). No more than 2% of motifs were allowed to pass the threshold in the ‘no-effect’ direction. All TFs for which the SNP created new best sites in caQTL versus no-effect enhancer above that threshold were also highly significant (all individual p-values < 0.00006) and would survive even a stringent ‘Bonferroni’ p-value adjustment. We repeated the analysis to look for ‘repressors’ by asking how often a caQTL versus no-effect SNP creates a new best site when going from the more to the less accessible genotype.

To compute a TF’s context score, we used a linear regression model by relating a TF’s top score in the less accessible genotype to the binary enhancer status (caQTL or no-effect) and the overall ATAC-seq coverage of the given enhancers for each TF separately:

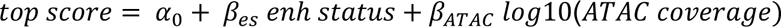

With α_3_ being a TF-specific intercept, and β_B7_ and β_FGFH_ the respective coefficients relating the top score to enhancer status and overall accessibility. Using ATAC-coverage removes the contribution of a TF to overall accessibility, which confounds the ‘caQTL’ versus ‘no-effect’ label. caQTLs are identified using count statistics, meaning that it is easier to detect a caQTL in highly accessible elements with high counts. Therefore, accessibility-driving TFs that are not specific for the creation of a ‘caQTL’ environment will wind up enriched in caQTL enhancers when accessibility is not controlled for. The entire procedure was repeated across the 50 different samples of ‘no-effect’ enhancers, meaning that each TF will have 50 slightly different β_B7_coefficients. To obtain a final TF context score, we extracted the average of the 50 t-statistics corresponding to each β_B7_. Positive t-statistics relate to TFs whose top binding sites score higher in caQTL than in no-effect enhancers. Importantly, to avoid confounding with the SNP itself, before fitting a model for a given TF, we dropped out all enhancers in which the SNP created a new best site for this particular TF, thus guaranteeing that the context score only captures the surrounding sequence, but not the SNP itself. We define a threshold of t > 3.2 to define context TFs, derived using the approach already described for the initiator score (defining a boundary based on no more than 2% of TF motifs falling within the ‘enriched in no-effect’ enhancers). In the case of context scores, the only motif significantly enriched in the context of no-effect caQTLs is CTCF, which is explained by its rather independent mode of action that is unlikely to require ‘cooperative’ RF clusters.

### Defining TF classes, overlap with USTs and TF association with enhancer annotations

TF classes were defined considering the accessibility score last. That is, if a TF was enriched at the SNP (log2-Odds Ratio > 0.9), it obtained the ‘-initiator’ label independent of whether it had a significant accessibility score or not. Next, initiator motifs were subdivided into context- or independent-initiators based on whether they scored high in the context score or not (context score t-statistic > or < 3.2). TFs whose binding sites were enriched in the context but not at the SNP were labeled as context-only TFs. Finally, TF motifs enriched for accessibility, but not covered in either of the two other scoring metrics were labeled USTs (accessibility score t-statistic >= 4). The name UST was chosen based on overlap statistics. For this, we took the annotation tables provided in the Supplement of the original Universal Stripe TF publication^46^ and computed the overlap between USTs and each motif class separately. Significance was determined using Fisher’s Exact Test.

To associate TFs with specific enhancer annotations, we retrieved regulatory enhancer annotations derived from DNA Hypersensitive Sites (DHS) Index^48^ across hundreds of cell types. We focused specifically on enhancers annotated as either ‘lymphoid-specific’ or ‘tissue-invariant’ to represent LCL-specific and universal functions of TFs. For each TF separately, we first split our random set of enhancers into two groups, those with above or below average top binding site strengths (centered around the mean top score for each TF). Next, we computed the ratio of ‘lymphoid-specific’ versus ‘tissue-invariant’ enhancer annotations for each of the two groups and took the ratio of ratios as an indication for a biased binding of the TF towards specific enhancers. For example, if a TF has a larger lymphoid-vs-invariant ratio in the above average binding site group, its binding is biased towards lymphoid-specific elements. Conversely, if the ratio is smaller, than TFs favor binding to tissue invariant enhancers.

### Allele-specific accessibility in GM12878

We performed Allele-Specific Accessibility (ASA) detection across the whole genome using ATAC-seq data downloaded from ENCODE^77^ (Experiment: ENCSR637XSC). In particular, we focused on the GM12878 sample, since we have a phased genotype available from the Genome in a Bottle database (https://www.nist.gov/programs-projects/genome-bottle). Then, we 1) aligned the *.fastq* file on the hg19 reference genome (GRCh37, release 75 from Ensembl) using bwa mem^72^ (v0.7.17-r1188), 2) marked duplicated reads using Picard^78^ (v2.17.8), and 3) counted the reads for each allele of the variants described in the phased genotype VCF file using Freebayes^79^ (v1.3.4) with the following options (--report-monomorphic, --only-use-input-alleles, --min-alternate-fraction 0, --variant-input [vcf file]). Of note, we discovered and reported a bug in Freebayes, which required to process each chromosome separately in order for the ASA calculation to finish properly. Finally, we summarized the ASA files using custom R script, focusing on heterozygous calls and counting the RO (reference) vs AO (alternate) allele calls.

### SuRE-seq library design

Starting from caQTL-probability and GC matched LOCAL and CM-Lead enhancers (see section below), we chose equal numbers from each category, prioritizing elements in the following way: caQTL SNPs that were heterozygous in the reference cell line GM12878, neighbors and dependent peaks that were annotated as ‘distal’ or ‘intronic’ and with similar distances from their respective caQTL enhancers. CaQTL elements were centered around the SNP and extended by 135bp on each side. Neighbors and dependents were centered at their respective peak centers, extending sequence up- and downstream by the same number of bps. The designed sequences were submitted to ANNOGEN (https://www.annogen.bio/) for SuRE^TM^ screening services using the target cell line GM12878. Transfections and data acquisition were handled by ANNOGEN, following the transfection protocol described here^80^.

### SuRE-seq data analysis

We processed SuRE-seq count tables as follows: we required a minimum barcode coverage of 2 from the paired-end sequencing reads generated for the barcode-to-element mapping (‘Reads’ in the raw count tables). All barcodes that mapped to more than one SuRE element were removed, as well as all elements that had less than 4 unique barcodes. Further, we required that the barcodes were observed at least once in the plasmid count columns. Since there is no relationship between plasmid and mRNA library prep, the replicate plasmid count columns were combined by adding them up for each element respectively. Processed tables for both ‘for’ and ‘rev’ element orientations were merged into a joint file, as orientations are not meaningful for enhancers and were simply defined by the genomic sequence of the ‘+’ strand.

Next, we computed the SuRE activity for each element-barcode combination as the log2 ratio of mRNA over plasmid DNA counts. We did this for both replicates separately, as well as for the combined replicate mRNA counts. The former was used to compute replicate agreement, and latter for all downstream analyses. To compute replicate agreement, each element was first summarized by its average SuRE activity across barcodes, including either orientation of the element, but keeping the genotypes separate, before computing the Pearson correlation coefficients between replicates.

sureQTLs are defined as caQTL elements whose average SuRE activity across all barcodes in one genotype differs significantly from that in the other (less and more accessible). Significance is assessed using Student’s t-Test and the ‘Bonferroni’ method to adjust the resulting p-values. The sureQTL - eQTL overlap is calculated by first extracting a caQTL’s probability to colocalize with an expressionQTL (eQTL; given in the variant table of the original LCL study^43^; see first Method section) for all caQTLs tested in SuRE. The empirical cumulative distribution function of the total set is compared to that of subsets, conditioning on either all sureQTLs, or sureQTLs with increasingly large effect sizes.

TF enrichment at sureQTL SNPs was computed as the log2 odds ratio of the frequency with which sureQTL and caQTL-only SNPs create new best binding sites for a given TF.

The relationship between SuRE activity and a TFs top binding site score was assessed by computing the Pearson correlation between the average SuRE activity in the less accessible genotype element and the binding site with the highest score within the 270bp defining the element (top scoring). For the SNP effect size comparison to ASA, elements were first split by their sureQTL status before computing the correlation between sureQTL effect (defined as the log2 difference in means between the more and less accessible genotype) and the log2 fold change in allele-specific ATAC-seq counts covering the SNP in the less and more accessible genotype in GM12878 respectively.

### Inferring combinations of SNP-affected and -neighboring top scores within caQTL elements

Analyses were performed using an ‘all-by-all’ approach. For each TF separately, the set of high-confidence caQTL elements was split based on whether the SNP created an enhancer-wide new best binding site or not. Comparing between the two splits, we iterated through all remaining TFs and computed the difference in distributions across top scores by performing a Student’s t-test. The difference in average top scores for each motif-by-motif combination is reported as the t-statistic, which corresponds to the color palette in Figure 3b and Supplemental Figure 3.

### STARR-seq library design and cloning

Three distinct 110 bp long random sequence embeddings were generated by uniform sampling with replacement across the four DNA bases (A, C, G, T). The optimal core motif sequences for 8 TFs (not-enriched but highly expressed: MYC = CCACGTGC, independent-initiator: NFKB/REL = GGGAAATTCCC, CTCF = CCACCAGGGGGCGC; context-initiators: SPI (PU1) = AAAGAGGAAGTGA, IRF = GAAAGCGAAACT, RUNX = TTTGTGGTTT; context-only: MEF2 = GCTAAAAATAGAA, FOX = CTGTTTACTTT) were retrieved from the HOCOMOCO^3^ database with an orientation chosen at random. For homotypic and heterotypic motif combinations, a 5-bp or 10-bp spacer was inserted between motifs, as well as added to the flanks of the first and last motif respectively. Each motif or motif-spacer-motif combination was placed within each of the three random sequence contexts such that the center of the insert aligned with the center of the random sequence. Each sequence was flanked by forward and reverse primer binding sites for subsequent library amplification and barcode addition. The in-silico designed oligos were ordered from Twist Biosciences.

Libraries were amplified with primers carrying a 12-bp random barcode (towards the 3-prime end), Nextera sequencing adapters, as well as overhangs for assembly into the STARR-seq vector (Addgene #99296; **Supplemental Table 1**) according to the manufacturer’s instruction (10 to 11 cycles of amplification). Amplified oligo pools were cloned into the STARR-seq vector by Gibson Assembly (NEB) and transformed into Dh5A high-efficiency competent cells (NEB). Transformed cells were spread on Ampicillin plates to achieve ∼50,000 individual clones (7-8 individual transformations) for each 5-bp and 10bp-spacer library (∼150 individual barcodes for the 186 (5-bp) or 149 (10-bp) motif combinations in 3 random sequence contexts).

Colonies were scraped from the Agar plates and transferred to liquid growth medium (Luria Broth) and grown at 37C for 2 h before plasmid DNA isolation (Maxiprep kit Invitrogen).

### STARR-seq transfection, RNA isolation and library preparation

MEC1 cells were cultured in IMDM media (Gibco) supplemented with 10% Fetal Bovine Serum and 1% Penicillin/Streptomycin to obtain ∼50 million cells. Transfections of 5-bp and 10-bp libraries were done in duplicates using ∼6*10^6^ cells and 30 *μ*g STARR-seq plasmid per replicate. Transfections were carried out with the Neon Transfection System (Thermo Fisher) as follows: 3 pulses of 20 ms at 1200 V in R buffer using 100 ul tips. Cells were harvested 24h post transfection and lysed in Trizol (Thermo Fisher). RNA was extracted by chloroform extraction, followed by isopropanol precipitation with a 1:1 ratio. RNA pellets were resuspended in RNAse-free water and digested with DNAseI for 20 min at room-temperature (Zymo). DNAseI-treated RNA was purified with a RNA-clean&concentrator-25 kit (Zymo). A total of 10-12 *μ*g RNA was reverse transcribed using the STARR-seq specific RT primer (**Supplementary table 1**). For each *μ*l of Maxima H minus RT (Thermo Fisher) 3 *μ*g input was used. RT reactions were diluted 1:4 with RNAse & DNAse-free water before further amplification. The splice-junction PCR was run using the STARR-seq thio-splice-junction primer and a custom reverse primer spanning the SalI restriction site used for linearizing the STARR-seq vector and the read1 Nextera adapter sequence inserted with the library (**Supplemental Table 1**). About one third of the RT reaction was amplified for 19-22 cycles. Illumina barcodes were added by a final PCR with 8 cycles using the Illumina Nextera guidelines. After each amplification step, PCR products were cleaned up using AMPure XP magnetic beads (Beckman Coulter). Plasmid input libraries were generated by amplifying the plasmid library directly with Nextera Read1 and Read2 indexing primers for 8 cycles (using a total of 600 ng plasmid in 6 separate PCR reactions). Plasmid libraries were sequenced on a MiSeq Illumina desktop sequencer at EPFL’s Gene expression Core Facility with a 150-cycle kit with the following read configurations: PE 80-8-80. RNA libraries were sequenced either on the same MiSeq run or separately on a NextSeq desktop sequencer at the same facility.

### STARR-seq data analysis

Paired-end sequencing data for the input plasmid libraries were used to assign random 12bp-barcodes to the different motif combinations in the three random sequence contexts using custom python and R scripts. Barcodes that mapped to more than one insert sequence were discarded from the downstream analysis. 12-bp barcodes were counted separately for both the plasmid DNA input and the two RNA replicates using a custom python script. Enhancer activity was computed by taking the log2 of the RNA to DNA ratio for each barcode and normalizing by the average activity of barcodes corresponding to the respective random-sequence contexts alone. As such each ‘motif-based’ activity represents activity above or below the expectation given by the random sequence context alone. To compute the average activity for a specific motif combination, the average log2 RNA/DNA ratios across all respective barcodes were computed first from which the average ‘random sequence context’ activity was subtracted in a subsequent step. For all analyses, only barcodes with at least 5 counts in the input plasmid library were considered.

To compute cooperativity for heterotypic two-motif combinations, two approaches were considered: Expressing cooperativity as the difference between the average ‘random-sequence-context-normalized’ heterotypic activity and i) the sum of the respective monomeric motif activities, or ii) the average of the two homotypic two-motif activities. For example, cooperativity induced by the combination of a ‘<MEF2-PU1-MEF2>’ site could either be computed by subtracting (2x <MEF2> + 1x <PU1>; i) or by (2/3x ‘<MEF2-MEF2-MEF2>’ + 1/3x ‘<PU1-PU1-PU1>; ii). The latter accounts for the effect two or more binding sites may have on enhancer activity independent of the motif identity (i.e., we observe a general downward trend in activity with each TF motif added). Cooperativity for trimeric or tetrameric motif combinations was calculated using the second approach.

### Computing TADs and low complexity domains within TF non-DBDs

Information on TF domains with start and end positions was extracted using the Ensemble based annotation package (EnsDb.Hsapiens.v86). To focus on non-DBDs and thus minimize the bias potentially introduced by DBD-similarities among members of the same TF family, we identified all protein domains associated with DNA-binding and removed them from the full-length sequence. To determine whether the non-DBD sequence contained a trans-activation domain (TADs), we used the approach described in Staller et al^51^. Briefly, for each TF we considered all 39 aa windows specific to the non-DBD and computed the total charge (counting up K and R and subtracting D and E amino acids) as well as the total number of W, F, Y, and L amino acids. TADs are then defined as domains that meet the following criteria:

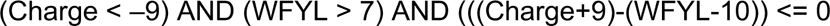

If a TF contained at least one 39aa window in its non-DBD that met the above criteria, we counted this TF as a TAD-TF. For each TF class, we then computed the contingency table of how many TFs were TAD-TFs or not, comparing to the group of unlabeled TFs. Significance was assessed using Fisher’s Exact Test.

To compute the fraction of amino acid windows with low complexity, we used the DBD-removed sequences and computed the number of unique aa within a 20 amino acid window as described in Van Mierlo et al^54^. If there were less or equal than 7 unique aa within a 20 aa window, we counted the window as a low complexity unit. The total low complexity fraction is then computed by dividing the number of windows where this criterion is met by the total number of windows. Note, we concatenated the amino acid sequences that were split by removing the DBD. At the breakpoint there is a non-existing amino acid stretch. However, since the goal is to compute low complexity which does not rely on specific folding, as is the case for TADs, we considered this to be the least problematic approach of several alternative choices: Keeping the DBD has obvious biases, as DBD families dictate motif groupings and have a significant sequence overlap from one family member to the next. Breaking the protein up into two or more stretches results in artificial end and start points, which disrupt the sliding window approach and leads to an undercounting of DBD-flanking amino acids.

Statistical significance between TF classes was assessed using a Mann-Whitney test.

### Association between TFs and epigenomic states

Fastq files for H3K27ac and BRD4 ChIP-seq experiments in GM12878 cells were retrieved from ENCODE (accession numbers: ENCFF000ASP, ENCFF000ASU for H3K27ac) or the SRA archive (SRR1636861 for BRD4). Raw fastq files were aligned to the hg19 reference genome using the bwa-mem^72^ algorithm. Alignment files were sorted, indexed, and de-duplicated with samtools^73^ and subsequently transformed to the ‘bigwig’ format using the bamCoverage command from deepTools^74^.

BRD4 or H3K27ac signal at a given enhancer was extracted as the cumulative signal within +/– 650 bp around the peak centers of all enhancers included in the random samples (see section about random enhancers). 650bp was chosen to allow for up to 2 additional nucleosomes surrounding the accessible region defined to +/– 350 bp around the peak center.

To compute the association of a TF with IP signal, we used a log-linear model, relating the cumulative IP-seq signal at a given enhancer to a TF’s top score. We used the scrambled top scores for the same TF as an additional covariate and fit the following model separately for each TF:

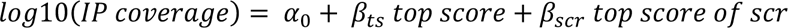

With α_3_, β_67_, β_7;<_ being respectively the TF-specific intercept, and the coefficients relating IP coverage around an enhancer to the TF top score in the actual or scrambled enhancer sequence. We extracted the t-statistic of β_67_as the evaluating metric.

Additionally, we fit separate models using the cumulative ATAC-seq coverage at the enhancer (+/– 350 bp around the peak center) as an additional covariate to account for the potential confounder of increased crosslinking independent of TF binding, but rather an artifact of the experimental assay:

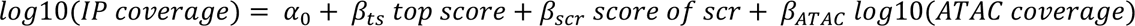

Similarly, we extracted the t-statistic of β_67_ as the evaluating metric for each TF.

### Defining independent and CM enhancer pairs

To define LOCAL and CM-leads, we started with the caQTL table defined in the first paragraph of the Methods, which simply required that the SNPs have a (‘P_Lead’ * ‘P_caQTL’) > 0.7 and to fall within 350 bp of the peak center, as well as the peaks to be annotated as ‘Distal’ or ‘Intron’ and have a minimum cumulative ATAC-seq coverage of 10 reads. Each caQTL enhancer was then annotated based on whether it was contained within the CM-annotation file (‘dag.txt’ for directed acyclic graph). Specifically, we labeled all caQTL enhancers as ‘CM-leads’ if they were contained in the first column of the ‘dag’-file. If a caQTL enhancer was not in any of the 5 columns of the dag file, they were labeled as LOCALs.

To generate matched controls between LOCALs and CM-leads, we considered GC content as well as the ‘potency’ of the SNP (‘P_caQTL’). The former is important to avoid biases in TF binding site scoring and the latter to guarantee that LOCALs have the same odds of revealing a change in a neighboring peak. LOCALs in general have lower P_caQTL values, which means there is a higher degree of uncertainty of whether a change in a neighbor is either absent or whether it simply cannot be detected because the initial change in the LOCAL is too small. Matching P_caQTL values resolves this uncertainty. Similar to the caQTL versus ‘no-effect’ enhancers, we generated 50 different subsamples of LOCALs matched in both their GC content and P_caQTL values to true CM-leads. Sampling was executed as previously described. In addition, we only retained caQTL enhancers if they had at least one neighboring peak or a respective ‘dependent’ that was also annotated as an ‘enhancer’. As CM-dependents, we considered only those peaks in the second column of the ‘dag’-file which represents a direct dependency hierarchy.

Since LOCALs can have two ‘enhancer’-neighbors and CM-leads can have more than one ‘enhancer’ CM-dependent, we also included a sampling step, which retains only one neighbor or CM-dependent per LOCAL or CM-lead. The choice is semi-random and executed in the following manner: if the neighbor or CM-dependent has a SNP heterozygous in GM12878, retain it. If both neighbors or several CM-dependents have a heterozygous SNP in GM12878, choose the one that has the highest coverage followed by the one closest to the peak center. This deterministic sampling is done to assess the cis-acting mechanism of CMs in GM12878. If none of the enhancer neighbors/CM-dependent has a heterozygous SNP in GM12878, choose one enhancer at random. This approach results in 50 samples of independent- and CM-enhancer pairings, which vary in their composition of included LOCALs, as well as different LOCAL-neighbors and CM-lead and -dependent pairs. Retaining only one CM-dependent is necessary, as we would otherwise count some CM-leads twice or multiple times, which can bias the TF binding site inference. The more dependents, the more potent a CM-lead, the more biased the TF composition, and the less generalizable the result.

In a separate sampling step, which is targeted specifically to assess TF binding site preferences in neighbors of CM-dependents, we find the closest LOCAL-neighbor distance match for each CM-lead – CM-dependent pair within each of the 50 samples. Doing so, we down sample the LOCAL-neighbor pairs, ending up with distance-matched independent- and CM-enhancer pairs with equal numbers of pairs in each subcategory.

For illustration purposes, representative samples are chosen based on the sample that is closest to an average significance metric, i.e., the sample whose p-value assessing the P_caQTL values or the peak center distances is closest to the average p-value across all samples. Significance is assessed using a Mann-Whitney Test for values obtained for LOCALs and CM-Leads or independent- or CM-pairs respectively.

### Quantification of peak interactions in Hi-C and Micro-C data

The Hi-C interaction matrix at 1kb resolution were retrieved from GEO (GSE63525; for GM12878). Raw fastq-files of Micro-C experiments in GM12878 (Micro-C 800M fastq files) were downloaded from the following link https://micro-c.readthedocs.io/en/latest/data_sets.html and aligned to the hg19 genome using the bwa-mem^72^ aligner. We used the *Pairix*^81^ and *cooler*^82^ tools *(as described in sections Pre-Alignment to Generating Contact Matrix* https://microc.readthedocs.io/en/latest/data_sets.html) to build balanced interaction maps at 500bp resolution. Peak pair interaction frequencies were quantified for Hi-C and Micro-C data by first extracting all bins that overlapped individual peaks before averaging across all possible interactions among the respective bins of a peak pair. For example, if peak1 and peak2 of a peak pair overlapped 2 bins each, we computed the average of the 4 resulting interaction scores.

The comparison of interaction frequencies between independent- and CM-enhancer pairs was done for all 50 sub samples matched by the caQTL enhancer P_caQTL and GC content, as well as by the distance between peak pairs. Statistical significance was assessed using the Mann-Whitney test.

### Evaluating TF binding site preferences among CM and independent enhancer pairs

To compare TF binding site preferences, we first generated a background expectation for TF top scores, i.e., how good of a score one would expect when sampling an accessible enhancer at random. Specifically, we computed the mean and the standard deviation across the top scores for each TF in the random enhancer sample. TF-specific mean and standard-deviation were then used to standardize (Z-score transformation) raw top scores, making it possible to compare across TFs. Next, we computed the average Z-score for a TF across enhancers belonging to one of the 4 categories (LOCALs, CM-leads, neighbors, CM-dependents). We did this for each of the 50 sub samples of matched CM- and independent enhancer pairs. A final average Z-score for a given TF in a given enhancer category was generated by averaging across the ‘averages’ obtained for each of the 50 sub samples.

To assess the average Z-scores in LOCALs and CM-leads, we used samples matched by GC content and P_caQTL, but not distance. For neighbors and CM-dependents, we additionally required distance matching to minimize bias that could arise through evolutionary-driven, distance-linked mechanisms.

To assess differences at the caQTL SNP between LOCALs and CM-leads, we used GC content and P_caQTL-matched enhancer sets and computed the frequency with which the SNP creates a new top site within LOCALs and CM-leads respectively. The final fraction was computed by averaging across the fractions of each of the 50 generated sub samples.

### Assessing BETi sensitivity among CM and independent enhancer pairs

Paired-end RNA-seq FASTQ files belonging to replicates of either JQ1 or DMSO-treated MEC1 cell (three replicates per treatment/vehicle), were retrieved from the SRA (SRR7815327; SRR7815329; SRR7815330; SRR7815331; SRR7815332; SRR7815333). Paired-end reads were aligned to the hg19 (GRCh37.75) reference genome using the STAR aligner^83^, sorted, indexed, and de-duplicated with samtools^73^ and loaded into R using the ‘Rsubread’ package^84^. The ‘DESeq2’ package^85^ was used for differential gene expression analysis, keeping genes with at least 10 counts across all 2x3 replicates.

CM-lead and LOCAL enhancer to gene-assignment was done by finding the closest TSS of all genes with an ensemble id using the ‘biomaRT’ R package. Annotation data for Super Enhancers and active B-cell enhancers were retrieved from the Super Enhancer Archive^60^ (SEA v3.0). Genomic coordinates were lifted to hg19 using the liftOver tool from the USC Genome browser. Enhancers annotated as ‘SE’ or ‘E’ (‘active enhancer’ in this manuscript) in one of the following cell types were considered for downstream analyses (‘B_cell’, ‘GM12878’, ‘Primary B-CLL cells’, ‘GM15850 LCLs (FRDA)’). Note, target gene annotations of SEs and active enhancers were taken directly from the SEA table.

The overlap between SEs/active enhancers and independent- or CM-enhancer pairs was determined in the following way: For active enhancers only the respective caQTL enhancer was considered (LOCAL, or CM-lead). For SEs, both enhancers of a pair (LOCAL and neighbor, or CM-lead and -dependent) had to fall within the boundaries of the SE. We defined overlap as a minimum of 200bp for each respective enhancer, for example, both CM-lead and -dependent had to separately overlap with an SE region.

When specific enhancers were annotated as both active enhancer and SE, SE was prioritized to avoid double counting enhancers. Computing overlaps was performed for each of the 50 GC content and ‘P_caqTL’-matched sets of LOCALs and CM-leads and their neighbors or CM-dependents. Note, the variability in log2 fold changes in expression seen in the CM-SE gene group is due to the fact that each sampling, albeit considering all CM-leads, can include a different CM-lead to CM-dependent assignment, as the latter is chosen at random from several potential dependents. In rare cases, this can also impact the set of the CM-active enhancers. We defined CM-active enhancers based on CM-leads only, however, the SE annotation supersedes the active enhancer annotation, which can lead to a CM-active enhancer either moving to the CM-SE group or not, depending on which dependent was chosen in the random sampling step.

Significance assessment comparing groups was done using the Mann-Whitney test.

## QUANTIFICATION AND STATISTICAL ANALYSIS

All statistical analyses were carried out in R. Raw p-values are visualized as . = 0.05 < p < 0.1, * = p < 0.05, ** p < 0.01, *** = p < 0.001, **** = p < 0.0001. When applicable, multiple testing correction was performed and is indicated in the respective **Methods** section.

