## Supplemental Material for "Context transcription factors establish cooperative environments and mediate enhancer communication"

**Supplemental Table 1 STARR-seq primer design**

| Name | sequence | description |
|---|---|---|
| STARR_lib ampli_12bp BC_rev | ctgctcgaagcggccggccgaattcgtcgacTCGTCG GCAGCGTCAGATGTGTATAAGAGACAG NNNN NNNN NNNN GCTGTCGGATCCGT | lower case = overlap with STARR-seq plasmid including AgeI restriction site;<br>blue/grey = Nextera i5 (Read1);<br>NNN = random barcode;<br>black = library fixed flank |
| STARR_lib ampli_for | cttgttgaattagattgatctagagcatgcaccggtGTCT CGTGGGCTCGGAGATGTGTATAAGAGACA GCGTGAGAGAACGCTC | lower case = overlap with STARR-seq plasmid including SalI restriction site;<br>red/grey = Nextera i7 (Read2);<br>black = library fixed flank |
| STARR_RT_primer | CTCATCAATGTATCTTATCATGTCTG | STARR RT primer |
| STARR_Spl-jct_for_thio | GTCGTGAGGCACTGGGCAGGEZEC | Splice-junction forward primer (EZE represents TGT but with phosphorothioate bonds) |
| STARR_Spl-jct_rev | CGACTCGTCGGCAGCGTCAG | Splice-unction PCR reverse primer |

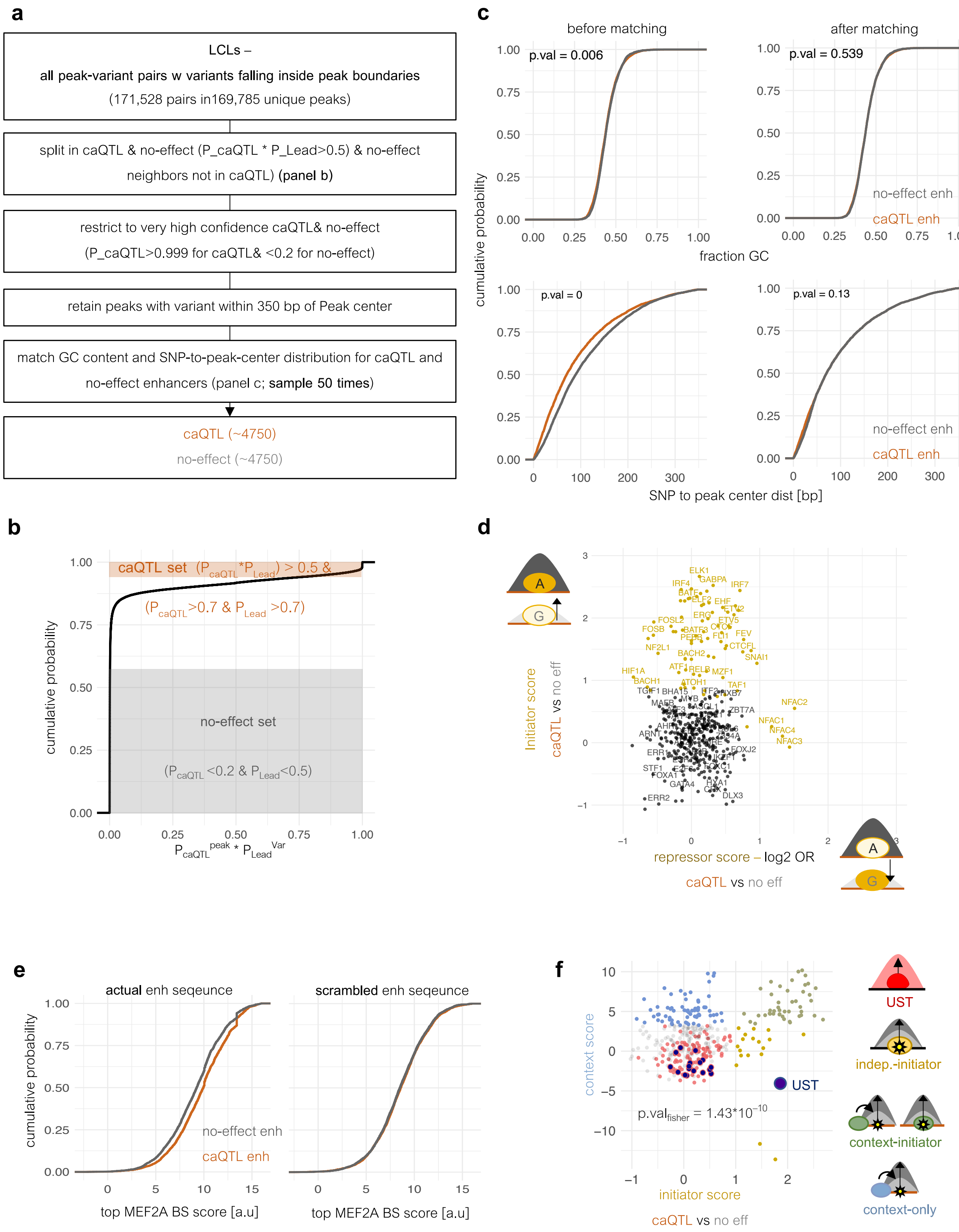


**Supplemental Figure 1 Data processing and characterizing TF function**

**a** Peak and variant data processing to obtain ‘no-effect’ and ‘caQTL’ enhancers. **b** Empirical cumulative distribution of a variant (SNP) to be both a caQTL and the ‘Lead’ variant indicating the thresholds used to define no-effect and caQTL sets. **c** Empirical cumulative distributions of GC content and the ‘SNP to peak center distance’ before and after matching no-effect to caQTL enhancers. A representative across 50 independent samples is shown for the no-effect enhancer set. **d** Log2 odds ratio of the number of SNPs creating new best binding sites for a given TF in either no-effect or caQTL enhancers (Fisher’s exact test). The y-axis shows gain (initiator score) and the x-axis loss (repressor score) of the best binding site when going from the less to the more accessible genotype. Yellow color indicates significance. **e** Example of assessing a TF’s context score in two groups of enhancers. Shown are the empirical cumulative distributions of raw MEF2 bindings site (BS) scores (top scoring) within the less accessible version of GC-content matched no-effect or caQTL enhancers (left). A shift indicates preferential binding to one or the other subset. Absence of a shift in scrambled enhancer sequences (right) confirms specificity for a TF independent of sequence composition. **f** Overlap of USTs with the group of TFs associated with high DNA accessibility (red) but neither context, nor initiator score (Fisher’s exact test). Dark blue points represent known USTs.

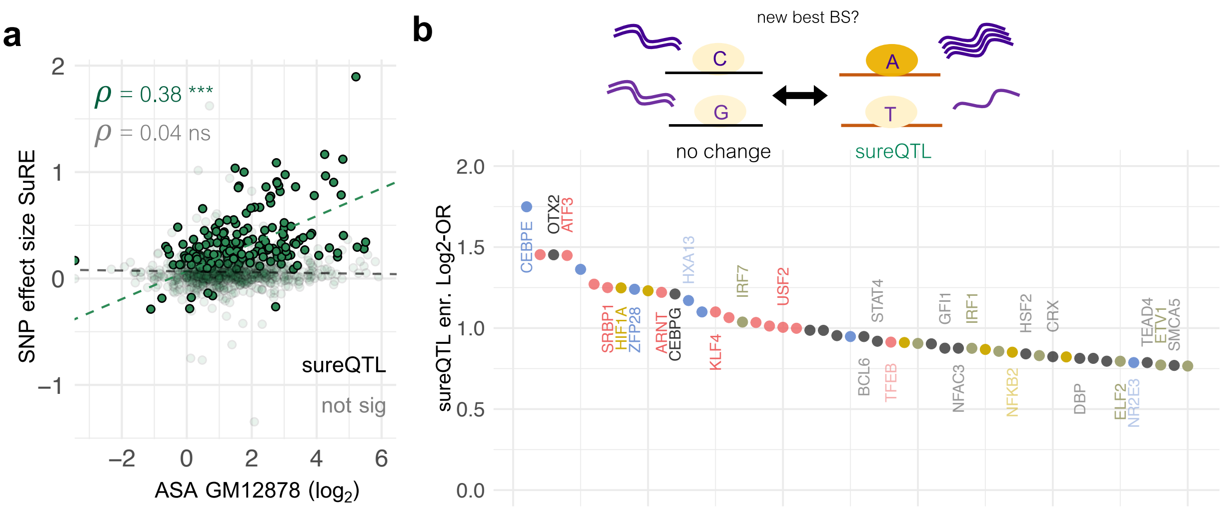


**Supplemental Figure 2 TFs associated with SuRE activity**

**a** SuRE SNP effect size expressed as the log2 ratio of the SuRE activity in the more versus the less accessible genotype of each fragment (y-axis) versus the allele-specific accessibility (ASA) change observed in GM12878 cells (x-axis). Bold and transparent colors differentiate sureQTLs from SNPs that are only caQTLs. Only sureQTLs show a correlation between the SuRE activity effect size and ASA. **b** 50 TFs with the largest log2 odds ratio of having a sureQTL create a new best binding site compared to the remaining caQTL SNPs that do not change activity. Colors represent TF classes and color shade indicates whether the corresponding uncorrected p-value was smaller of equal to 0.01.

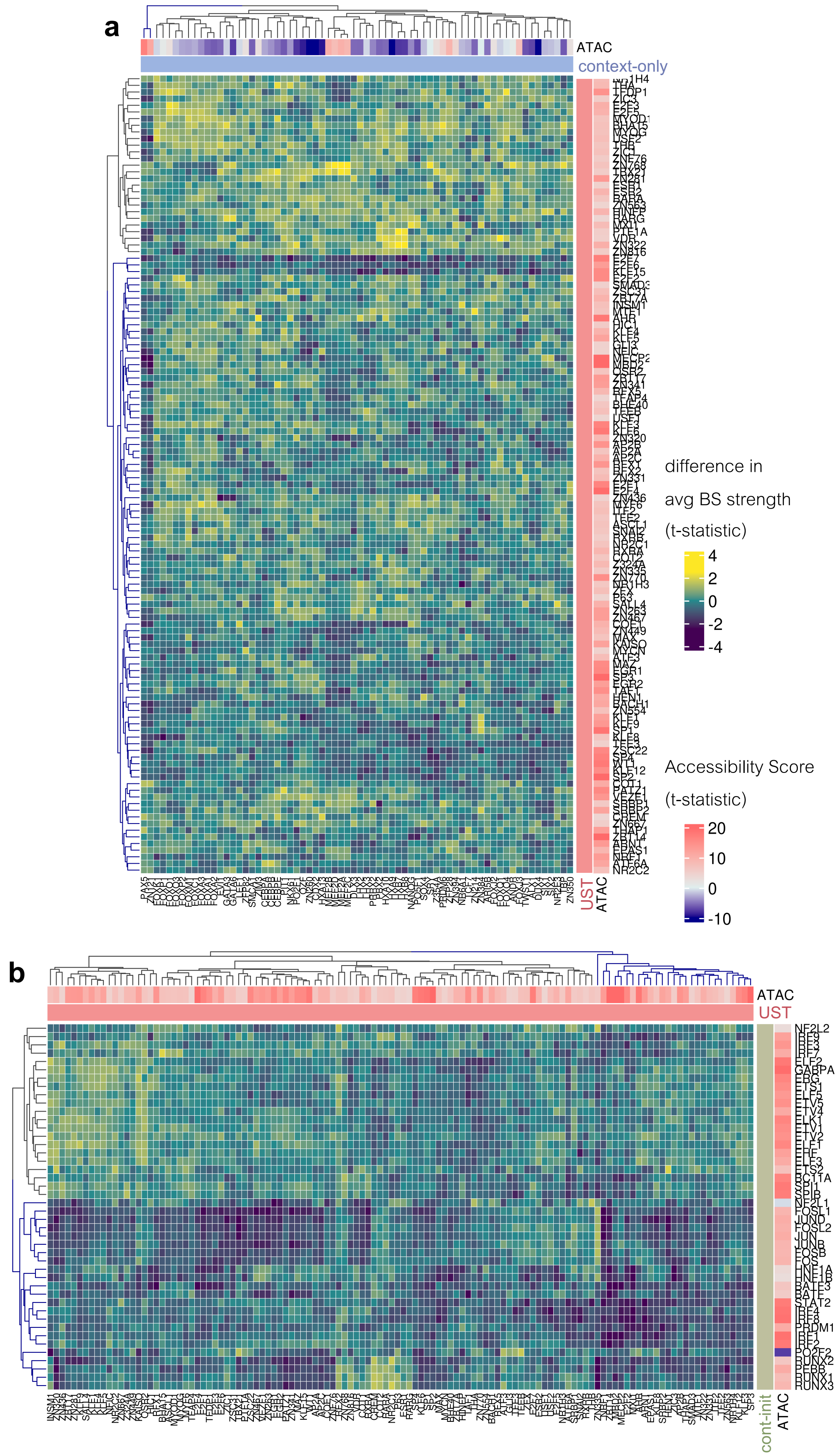
a
ATAC
context-only
UST
ATAC
difference in
avg BS strength
(t-statistic)
4
2
0
-2
-4
Accessibility Score
(t-statistic)
20
10
0
-10
b
ATAC
UST
cont-init
ATAC

**Supplemental Figure 3 Little evidence of specific pairings involving non-context TF pairs in caQTL enhancers**

**a/b** Hierarchical clustering based on the t-statistic computed for each TF pair as described in **Fig. 3a** and split in 2 main clusters (dendrogram colors). Columns contain context-only or USTs and rows USTs or context-initiator TFs respectively. The TF accessibility score is shown in the outermost annotation panel (blue-red color scale).

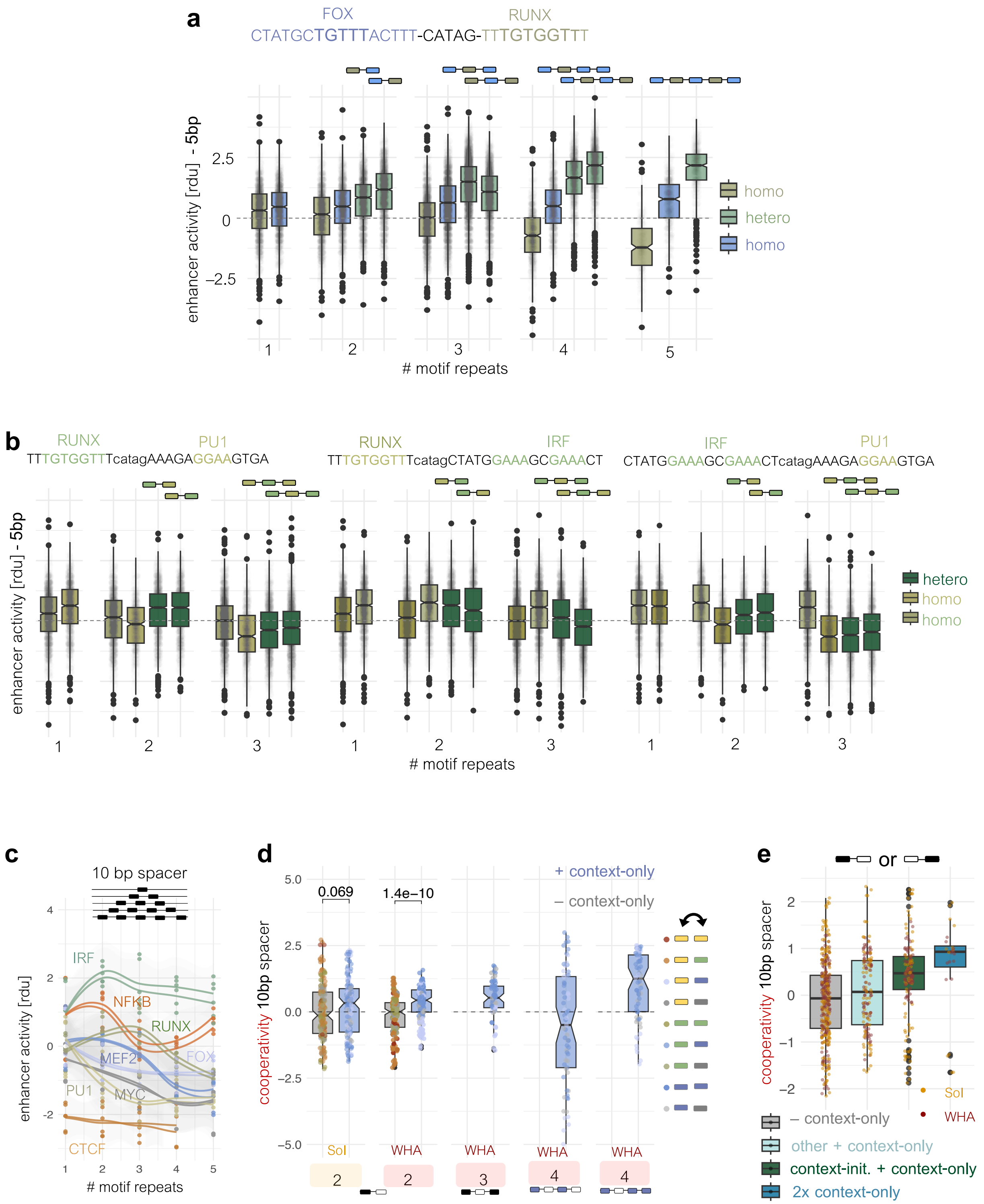

**Supplemental Figure 4 Testing for motif cooperativity using STARR-seq**

**a** STARR-seq activity for individual barcodes of homotypic or heterotypic motif combinations of RUNX and FOX motifs normalized by the average STARR-seq activity of the respective random sequence contexts alone (rdu = in terms of random units; y-axis). Motif orientation and 5bp-spacer sequence is given on top. Motif repeats are indicated below and the exact order of heterotypic repeats is indicated above each box. Colors indicate the corresponding TF motif (blue = FOX; yellow green = RUNX, blue-green = FOX & RUNX). **b** same as **a**, but for combinations among context-initiator motifs. **c** STARR-seq activity for homotypic motif repeats from the 10bp spacer library measured as the average across all barcodes and normalized using the respective motif-free random sequences as a reference (rdu = in terms of random units). Colors reflect TF classes. Lines are separate replicates and points belong to one of the three random sequence contexts. **d** Cooperativity for the 10bp spacer library (heterotypic-homotypic) (y-axis) as a function of motif number (x-axis schematics). Method indicated by SOI = sum of individual homotypic motifs or WHA = weighted homotypic average of same length. Colors indicate whether a motif combination contains a context-only TF or not (blue/grey). Color of points indicates different combinations of specific motif classes. Different sequence contexts and replicates represent individual points. **e** Same as **d** but focusing only on two motif combinations and splitting based on distinct motif class combinations (x-axis). (no context-only motif = grey; context-only & independent-initiator or unlabeled = cyan; context-only and context-initiator = dark green; 2 context-only = blue). Points are colored based on the methods used to assess cooperativity (cf. **d**).

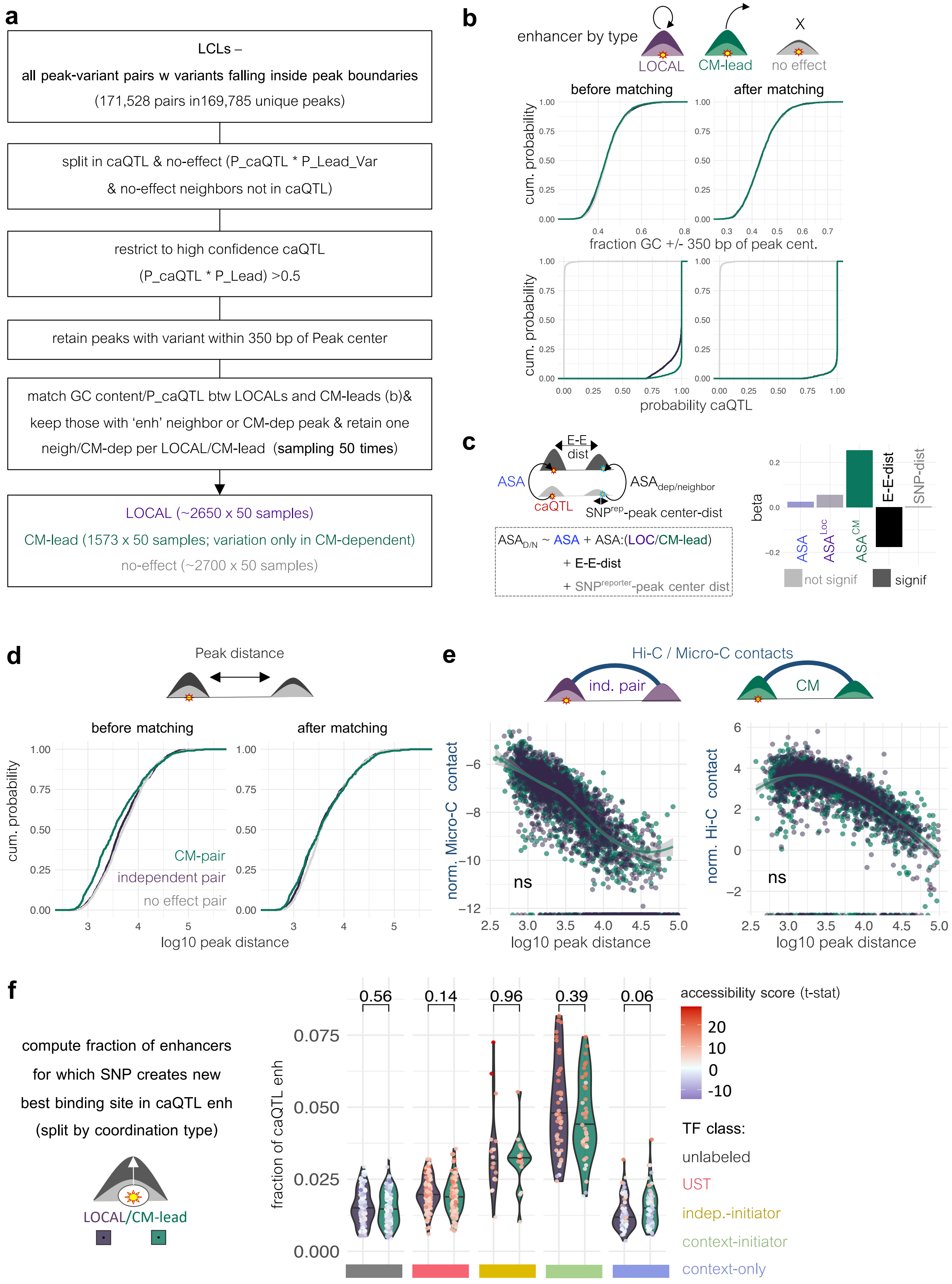

a
LCLs –
all peak-variant pairs w variants falling inside peak boundaries
(171,528 pairs in169,785 unique peaks)
split in caQTL & no-effect (P_caQTL * P_Lead_Var
& no-effect neighbors not in caQTL)
restrict to high confidence caQTL
(P_caQTL * P_Lead) >0.5
retain peaks with variant within 350 bp of Peak center
match GC content/P_caQTL btw LOCALs and CM-leads (b)&
keep those with 'enh' neighbor or CM-dep peak & retain one
neigh/CM-dep per LOCAL/CM-lead (sampling 50 times)
LOCAL (~2650 x 50 samples)
CM-lead (1573 x 50 samples; variation only in CM-dependent)
no-effect (~2700 x 50 samples)
b
enhancer by type
LOCAL
CM-lead
no effect
before matching
after matching
cum. probability
fraction GC +/- 350 bp of peak cent.
probability caQTL
c
E-E dist
ASA
ASA dep/neighbor
caQTL
SNP rep-peak center-dist
ASA D/N ~ ASA + ASA:(LOC/CM-lead)
+ E-E-dist
+ SNP reporter-peak center dist
beta
E-E-dist
SNP-dist
not signif
signif
d
Peak distance
before matching
after matching
cum. probability
CM-pair
independent pair
no effect pair
log10 peak distance
e
Hi-C / Micro-C contacts
ind. pair
CM
norm. Micro-C contact
norm. Hi-C contact
ns
log10 peak distance
f
compute fraction of enhancers
for which SNP creates new
best binding site in caQTL enh
(split by coordination type)
LOCAL/CM-lead
fraction of caQTL enh
0.56
0.14
0.96
0.39
0.06
accessibility score (t-stat)
TF class:
unlabeled
UST
indep.-initiator
context-initiator
context-only

**Supplemental Figure 5 Creating matched sets of communicating and independent enhancer pairs**

**a** Procedure to generate matched LOCAL and CM-lead enhancers. **b** Empirical cumulative distribution of GC content and P_caQTL values across different enhancer types before and after matching. Shown is a representative sample for LOCAL and no-effect enhancers **c** Assessing the *cis*-acting enhancer to enhancer communication within CMs in GM12878. Allele-specific accessibility (ASA) measured going from the less to the more accessible genotype at a reporter SNP within a neighbor or CM-dependent peak (effect size orientation is defined based on the lead variant using phased genome information) is modeled as a function of the ASA at either the LOCAL/CM-lead, the ASA conditioned on the enhancer type ($ASA^{LOC}$ for LOCALs; $ASA^{CM}$ for CM-leads), the distance between paired peaks (E-E-dist) and the distance between reporter SNP and peak center (SNP-dist). Plotted are the inferred coefficients (beta) of a log-linear model from a representative sampling of LOCALs. Enhancer-enhancer distance is negatively, whereas ASA at the CM-lead is positively associated with the inherited ASA at the (neighbor)/CM-dependent. Transparent bar color indicates non-significant estimates. **d** Empirical cumulative distribution of peak center distances split by enhancer pair type before and after distance matching. Shown is a representative sample from a total of 50 matched sets. **e** Peak-pair contact frequencies of distance-matched independent and CM enhancer pairs as measured by Micro-Capture-C or Hi-C (500bp and 1kb resolution respectively) in GM12878. Shown is a representative sample. **f** Fraction of SNPs creating a new best binding site for a given TF when going from the less to the more accessible genotype in either LOCALs or CM-leads. TFs are split based on TF classes.

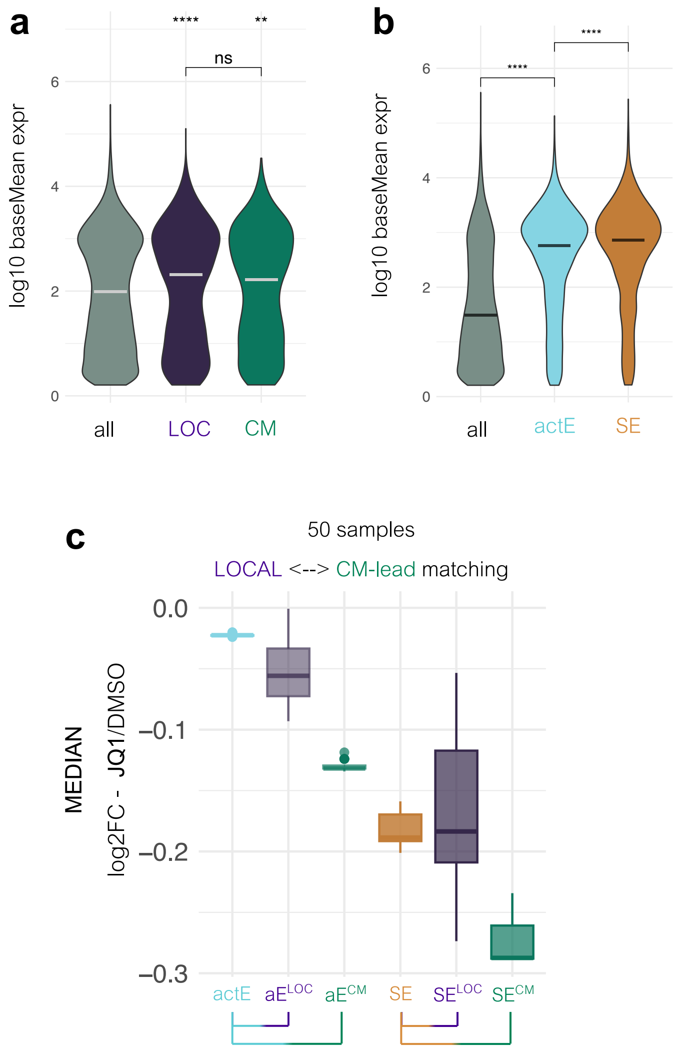


**Supplemental Figure 6 Comparison of LOCAL, CM and SE sensitivity to BETi**

**a** Mean expression levels of genes linked to either any, LOCAL or CM-lead enhancers. Enhancer to gene matching is based on the closest TSS. **b** Same as in **a** but using active B-cell enhancer (actE) and Super Enhancer (SE) annotations from SEA. Enhancer type and enhancer-to-gene mappings are taken from SEA. **c** Median log2 fold change in gene expression upon JQ1 treatment when subcategorizing active enhancers and SEs based on their enhancer communication status ($aE^{LOC}$ and $aE^{CM}$ for active enhancers overlapping LOCALs and CM-leads respectively; $SE^{LOC}$ and $SE^{CM}$ for SEs overlapping LOCALs + neighbors and CM-leads + CM-dependents respectively). Medians are shown for all 50 subsamples of LOCALs and CM-leads and their neighbors and CM-dependents.
